# REDD1 Regulates MERCS, Protein Synthesis and NMJ Stability in Fast Myofibers During Dexamethasone-Induced Muscle Wasting

**DOI:** 10.64898/2026.09.07.749905

**Authors:** D. Pereira, F. Simon, M. Di Gallo, E. Jauliac, G. Macaux, Z. Cepella, S. Polat, A. Campagna, O. Arnould, C. Bertrand-Gaday, S. Backer, A.H. Sinclair, S. Jacques, B. Izaac, A. Schmitt, P. Delobel, A. F. Pagano, E. Hucteau, J. Mallard, J. Rieusset, T. Rousseaux, B. Chabi, V. Ollendorff, G. Py, F. A. Britto

## Abstract

**Background:** Glucocorticoids cause skeletal muscle atrophy preferentially affecting fast glycolytic fibers, but the mechanisms involved in this fiber selectivity is unclear. REDD1 is a glucocorticoid-induced stress protein that limits muscle protein synthesis inducing atrophy. However, it remains unknown whether REDD1 exerts myofiber type-specific effects and through which precise mechanisms it regulates protein synthesis. We investigated the role of myofiber REDD1 expression in dexamethasone (DEX)-induced muscle atrophy, with a particular focus on its involvement in mitochondria–ER contact sites (MERCS), protein synthesis, and neuromuscular junction (NMJ) integrity.

**Methods:** We generated tamoxifen-inducible, muscle-specific REDD1 knockout mice (REDD1^fl/fl^HSA-CreERT2) and compared them with floxed littermates (WT) in a 2×2 design (WT/KO × PBS/DEX, 7 days). We combined single-nucleus RNA sequencing, RNAscope, immunofluorescence, transmission electron microscopy, proximity ligation assay, SUnSET puromycin labelling, western blot and RT-qPCR, and AdenoFATE1-mediated MERCS disruption in C2C12 myotubes.

**Results:** Glucocorticoid receptor and REDD1 transcripts were co-enriched in fast glycolytic fibers mostly atrophied by DEX (∼20%). REDD1 deletion in myofiber drove to lower basal muscle mass and fast fiber volume but protected them from DEX-induced atrophy. DEX inhibited protein synthesis (∼70%) in WT mice with no matching change in Akt/mTOR-pathway activity. In REDD1 KO mice, protein synthesis was already low and was not affected by DEX. DEX-induced REDD1 expression remodelled mitochondrial network and MERCS in a subcellular compartment-specific manner. The intermyofibrillar MERCS minimum distance shortened in both genotypes reaching pathological distances only in WT mice (WT ∼28→∼5 nm; KO ∼25→∼15 nm). Perinuclear MERCS and mitochondria-nuclei distances increased in WT mice only (∼18→∼45 nm and ∼130→∼460 nm). In WT mice only, DEX-induced alteration of the perinuclear mitochondrial network was associated with a loss of myonuclei accumulating mt-RNA and exhibiting an anabolic transcriptomic signature notably enriched in sarcomeric transcripts. These findings suggest that REDD1-dependent MERCS remodelling may regulate muscle anabolism beyond the control of mRNA translation, by shaping the myonuclear transcriptome. Finally, REDD1 localised to the NMJ and reduced endplate area during DEX treatment. Interestingly, MERCS were denser in NMJ than in myofiber body and we showed in vitro that FATE1-mediated MERCS disruption was sufficient to reduce protein synthesis and agrin-induced acetylcholine-receptor clustering demonstrating that REDD1 and MERCS are important for NMJ stabilization.

**Conclusions:** Muscle REDD1 links the glucocorticoid response to compartment-specific mitochondrial network remodelling, protein synthesis as well as NMJ stability in fast glycolytic fibers. Our results also show that REDD1 is important for maintaining basal mitochondrial network and protein synthesis homeostasis.

## Introduction

Chronic glucocorticoids treatments are widely used in clinic for their anti-inflammatory effects causing skeletal muscle wasting characterized by glycolytic fast fiber specific atrophy (1,2) . Because muscle mass is a determinant of mobility, metabolic health and survival, understanding glucocorticoid-induced atrophy is a key challenge for improving patient’s quality of life.

Glucocorticoids notably promote muscle wasting by dysregulating Akt/mTORC1-dependent protein balance (3). In addition, glucocorticoids treatment alters mitochondrial function (4,5). Mitochondria are the main source of ATP in the cell (6). This energy is required for protein synthesis, one of the most ATP-consuming processes in the cell (7,8). Mitochondrial activity depends on the physical contacts between mitochondria and the endoplasmic reticulum (ER), called MERCS (Mitochondria-ER Contacts Sites). These contacts mediate the transfer of Ca^2+^ from ER to the mitochondria that is required for ATP production (9–11). A link between these contacts and protein synthesis has been reported in HeLa cells, in which an experimental alteration of MERCS was accompanied by a decrease in global protein synthesis (12). However, whether and how MERCS regulate protein synthesis in skeletal muscle has not been studied. The neuromuscular junction (NMJ) is also destabilised by glucocorticoids (13). The NMJ is continuously remodelled (14), and its proteins are renewed (15). Mitochondria are enriched on both sides of the synapse (16), and their age-related decline correlates with denervation (17). Together, these features make the NMJ a structure in which a limitation of protein synthesis and mitochondrial function could be particularly damaging, although this has not been examined in detail. In addition, whether MERCS are present and functional on the muscle side of the NMJ has never been described.

Regulated in Development and DNA Damage 1 (REDD1) is a stress-responsive protein ubiquitously expressed at low levels under basal conditions and upregulated by various stresses such as glucocorticoid exposure (18). We previously reported that REDD1 total and constitutive deletion preserves muscle mass in dexamethasone(DEX)-treated mice, indicating that REDD1 is required for glucocorticoid-induced atrophy (19). However, the expression of REDD1 across fiber types or NMJ and its specific role in glycolytic fibers atrophy or NMJ destabilization have never been described. REDD1 is known as a repressor of the master regulator of protein synthesis mTORC1, and its expression limits protein synthesis in skeletal muscle (19,20). Beyond this inhibition of the Akt/mTOR pathway, REDD1 has also been shown to bind GRP75, a component of the VDAC1–GRP75–IP3R complex, to disrupt this tether protein complex, disrupting MERCS (21).

In this study, we describe the role of REDD1 in glycolytic fibers atrophy and NMJ destabilization in response to glucocorticoid treatment. We notably focus on REDD1 effect on the integrity of MERCS and the control of protein synthesis. Establishing such a relationship would place inter-organelle contact sites among the determinants of protein synthesis in skeletal muscle, and extend this question to the neuromuscular junction, and glucocorticoid treatment where it has not been considered.

## Material and Methods

### Animals

All procedures followed EU Directive 2010/63/EU and were approved by the Languedoc-Roussillon ethics committee and by the French Ministry of Higher Education, Research and Innovation (APAFIS #46107-2023112815379359). Inducible skeletal muscle-specific REDD1 knockout mice were generated by crossing REDD1-floxed mice, kindly provided by the laboratory of A.H. Sinclair (22), with tamoxifen-inducible, muscle-specific HSA-CreERT2 mice.

Mice were maintained on a C57BL6/N background. Only female mice were used, consistent with previous studies investigating the role of REDD1 in glucocorticoid-induced muscle atrophy (19). Mice were housed in a conventional facility at 24°C under a 12h light / 12h dark cycle, with ad libitum food and water. To induce muscle-specific REDD1 deletion, mice were injected intraperitoneally (IP) with tamoxifen (MP Biomedicals, #156738, 20mg/kg/day) for 5 consecutive days. Control mice received tamoxifen every day. Dexamethasone (D2915 Sigma-Aldrich) was injected daily by IP at 200μg/kg for 7 days to induce atrophy. On the final day, a single 1mg/kg injection was administered and tissues were collected 5h later corresponding to the peak of REDD1 expression (19). Combining these two protocols allowed us to capture both the atrophic response and maximal REDD1 induction in the same animals, thereby reducing the number of mice used. Control mice received IP injections of PBS over the same period.

### Immunofluorescence

Immunofluorescence for fiber cross-sectional area and fiber-type composition analysis were realized using the FLASH method developed in our team (23). To label the NMJ and nuclei, isolated fibers of Tibialis Anterior fixed in 4% paraformaldehyde (PFA) were incubated with α-bungarotoxin conjugated to Alexa Fluor 488 (1:1000, #B13422 Thermo Fisher) and DAPI for 20 min at RT. Images were obtained using a spinning disk confocal microscope (IXplore model, Olympus manufacturer) equipped with a 60× objective and analysed using Image J software. NMJ morphology were analysed using NMJ-moprh macro (24).

### Proximity Ligation Assay and NMJ labelling

Proximity Ligation Assay (PLA) was performed on Tibialis Anterior isolated muscle fibers fixed in 4% PFA using the Duolink in Situ kit (Sigma Aldrich). After adhesion, fibers were permeabilized with 1% Triton X-100 in PBS for 1h at RT. PLA was then carried out according to the manufacturer’s instructions. Briefly, fibers were incubated with primary antibodies against VDAC (Mouse, Abcam #AB14734, 1 :200) and IP3R (Rabbit, Abcam #AB5804, 1 :200) overnight at 4°C, followed by incubation with anti-rabbit PLUS (DUO92002) and anti-mouse MINUS (DUO92004) PLA probes, ligation, and amplification with Far Red detection reagents (DUO92013). To label the NMJ and nuclei, fibers were then incubated with α-bungarotoxin conjugated to Alexa Fluor 488 (1:1000, #B13422 Thermo Fisher) and DAPI for 20 min at RT. Images were obtained using a spinning disk confocal microscope (IXplore model, Olympus manufacturer) equipped with a 60× objective and analysed using Image J software.

### RNA In Situ Hybridization

RNAscope was performed on isolated muscle fibers of tibialis anterior (fast fibers) or soleus (slow fibers) muscles using the RNAscope Multiplex Fluorescent Reagent Kit v2 (Catalog #323100, Bio-Techne) according to the manufacturer’s instructions. To assess REDD1 expression across fiber types, a probe for REDD1 (#493611, C1) was combined with intronic (pre-mRNA) probes detecting Myh4 (#539391-C2). In separate experiments, the REDD1 probe was combined with either Etv5 (#316961-C3). Channels were developed with HRP-based detection and revealed with Akoya Opal fluorophores. Nuclei were stained with DAPI (provided in the kit).

### Western Blotting and RT-qPCR

Western Blot and RT-qPCR were performed in quadriceps muscle protein samples with the same methods described in our previous studies (19,21). Cultured C2C12 myotubes were lysed in RIPA buffer (Sigma #R0278) supplemented with protease inhibitors (Sigma #P8340). Antibodies and primers used are provided in Table 1 and 2.

### Transmission electron microscopy

Immediately after dissection, the TA muscle from a separate cohort of animals, subjected to the same treatment protocol, was fixed in 2.5% glutaraldehyde and 4% PFA for 1 h at room temperature. The fixative was then replaced with fresh solution, and samples were kept at 4°C for 24h. Imaging was performed on a JEOL 1011 TEM with a Gatan Orius 1000 CCD camera.

### Quantification of MERCS and mitochondria morphology

Mitochondria–reticulum contacts were quantified using a Fiji macro as previously described (25). Mitochondrial morphology was analysed with AMITO, which provided mitochondrial diameter, shape, circularity, and minimum and maximum Feret diameters (26).

### Single nucleus RNA sequencing

For each condition, gastrocnemius from 3 to 5 animals were pooled. Nuclei were isolated using the nuclei extraction protocol of Miltenyi Biotec (130-128-024). Then, the nuclei were purified using the Anti-Nucleus Microbeads protocol (130-132-997). After that, nuclei were fixed 20 hours at 4°C using Fixation of Cells & Nuclei for GEM-X Flex Gene Expression of 10X Genomics (CG000782). Single-nuclei gel bead-in-emulsions (GEMs) were produced using a Chromium iX (10X Genomics). Sequencing libraries were created using the Chromium Single Cell FLEX Reagent Kit v2 (10X Genomics). We used NS 2000 system (Illumina, San Diego, CA, USA) to produce sequencing data. Fastq files were processed to make transcript count tables using Cell Ranger Software Suite 9.0.1 (10X Genomics). Downstream single-nucleus RNA-seq analysis was performed using Cell-Hub (27).

### C2C12 culture and treatments

The mouse myoblast cell line C2C12 was obtained from the American Type Culture Collection (ATCC, Manassas, VA, USA). C2C12 myoblasts were maintained at 37°C in a humidified atmosphere containing 5% CO2. Cells were propagated in growth medium (GM) containing high-glucose DMEM (D5796, Sigma-Aldrich), 10% fetal bovine serum, and 1% penicillin-streptomycin, in T75 flasks. For differentiation, cells were cultured in medium containing high-glucose DMEM, 2% horse serum and 0.1 penicillin-streptomycin. For the low-glucose experiment, cells were propagated in high-glucose DMEM as described above and switched at day 4 of differentiation, 16h before transduction, to glucose-free DMEM (11966025, Gibco, Thermo Fisher Scientific) supplemented with 5.5 mM glucose. At day 4 of differentiation, C2C12 myotubes were transfected with an adenovirus overexpressing FATE1 (Ad-FATE1) or a control adenovirus (Ad-GFP) kindly provided by Dr. Jennifer Rieusset (CarMeN laboratory, Inserm, Lyon, France), for 48h. During the last 16h, recombinant agrin (#550-AG, R&D Systems) was added directly to the medium at a final concentration of 1ng/mL to induce acetylcholine receptor (AChR) clustering (28,29). To measure protein synthesis, myotubes were incubated with [9,18uM] of puromycin for 30 min before lysis, and puromycin incorporation was assessed by Western blotting (SUnSET method).

### Statistical analysis

Statistical analyses were performed using GraphPad Prism 8.00 (GraphPad Software, San Diego, CA, USA), and data are reported as means ± SE. To examine the significance of observed differences between groups, we performed a two-way ANOVA, which allows the assessment of both main and interaction effects of two independent factors. Prior to analysis, data were tested for normality to ensure that the assumptions of ANOVA were satisfied. All statistical analyses were conducted using the significance threshold as follow: * p < 0.05; ** p<0.01; *** p<0.001.

## Results

### REDD1 deletion specifically in myofiber protects from glucocorticoid-induced fast glycolytic fibers atrophy

DEX increased REDD1 protein and mRNA levels in the skeletal muscle of WT mice (∼2 fold and ∼16 fold, Fig 1A, B and C). In REDD1 knockout (KO) muscle, REDD1 protein was largely abolished, the residual signal detected in KO DEX is consistent with expression in non-myofibers cells present in whole-muscle lysates (Fig. 1A, B, C and D).

**Figure 1.**
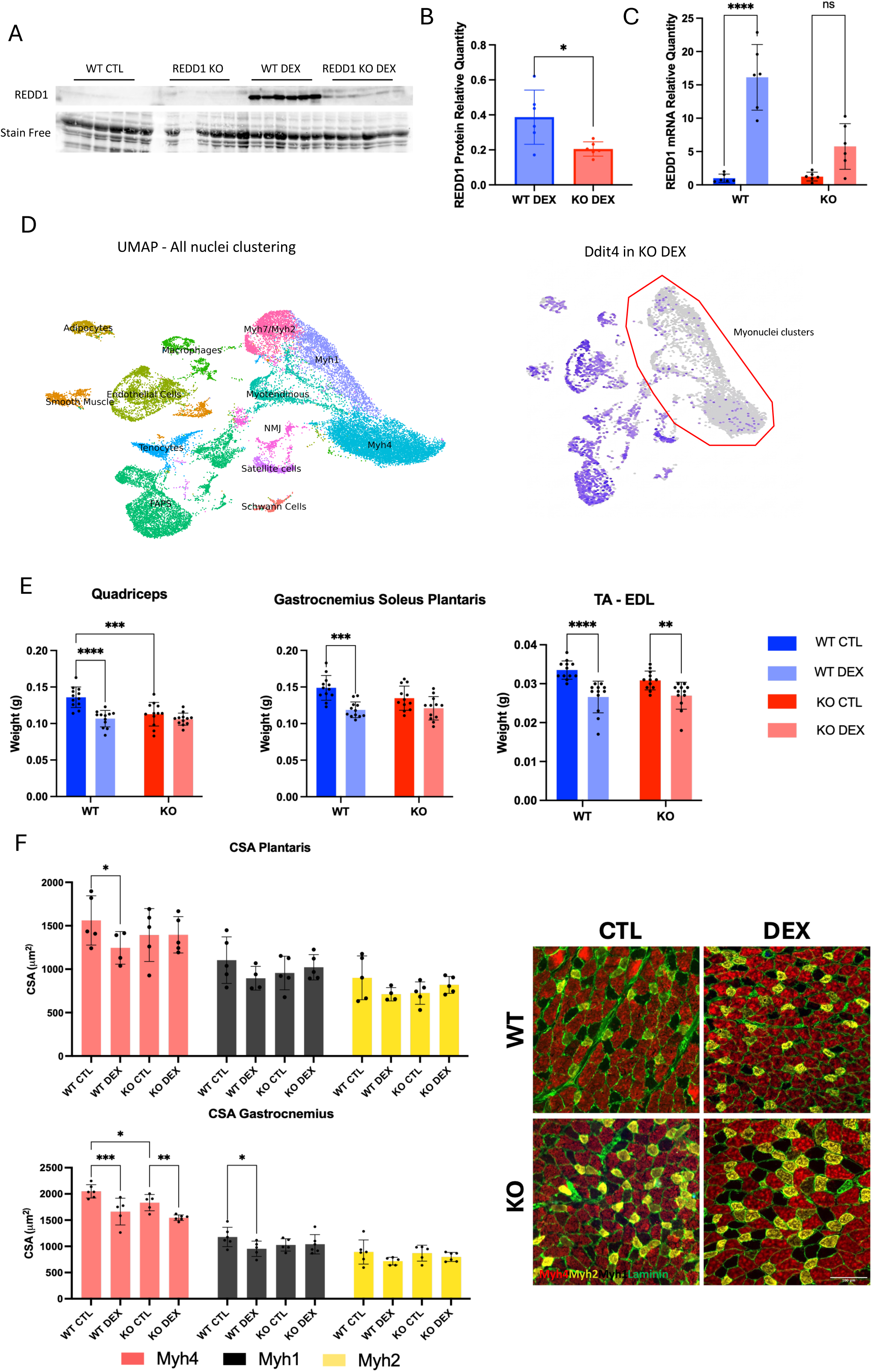
REDD1 is induced by dexamethasone in skeletal muscle and its muscle-specific deletion attenuates glucocorticoid-induced muscle atrophy. **A**. Representative western blot of REDD1 and total protein (Stain-Free) in quadriceps muscle of WT and musREDD1 KO mice treated with vehicle (CTL) or dexamethasone (DEX). **B.** Quantification of REDD1 protein normalized to Stain-Free in DEX-treated mice (WT DEX *vs* KO DEX). n = 6 per group; unpaired t-test *p = 0.0194. **C.** *Ddit4* (REDD1) mRNA relative quantity measured by RT-qPCR in WT and KO mice under CTL and DEX. Tukey’s post-hoc ****p < 0.0001; ns, not significant. n = 6 mice per group; two-way ANOVA followed by Tukey’s multiple-comparisons test. Gene expression was normalized to Rps9. **D.** Single-nucleus RNA-seq gastrocnemius muscle. Left: UMAP projection of all nuclei showing the major muscle cell population. Right: Ddit4 (REDD1) transcript distribution in KO DEX muscle, with the myonuclear clusters outlined. **E.** Muscle weights (g) of quadriceps, gastrocnemius-soleus-plantaris, and TA-EDL across the four groups. n=6; two-way ANOVA, Tukey’s post hoc. F. Fiber cross-sectional area (CSA, µm²) quantified by fiber type (Myh4, Myh1, Myh2) in plantaris and gastrocnemius; two-way ANOVA per fiber type, Fisher’s LSD post-hoc. Representative immunofluorescence of muscle cross-sections showing Myh4 in red, Myh2 in yellow, Myh1 unstained (negative fibers), and laminin in green.

We next characterized the baseline metabolic phenotype of the KO mice by indirect calorimetry. Under standard conditions, REDD1 KO mice displayed a higher respiratory exchange ratio (RER) than REDD1 floxed during light and dark phases (Fig. S1A), accompanied with a higher oxygen consumption (Fig. S1B). Both floxed genotypes showed a lower RER than the C57B6/N reference (Fig. S1A), suggesting that muscle specified REDD1 deletion was associated with an increase in whole-body energy expenditure.

DEX reduced mass of all muscles in WT mice while KO mice displayed a significant DEX-induced atrophy only in TA-EDL groups (∼-8%) (Fig. 1E). Quadriceps mass was already lower in KO mice in comparison to WT mice indicating a baseline reduction in muscle mass in the absence of REDD1 (Fig. 1E). DEX promoted reduction of Myh4 (type IIB, fast glycolytic) fibers CSA in both the plantaris and gastrocnemius (∼20%) in WT mice only (Fig. 1F). Together, these results show that REDD1 loss is associated with a modest baseline atrophy of fast fibers and protects glycolytic fibers from dexamethasone treatment.

### REDD1 and the glucocorticoid receptor are enriched in fast glycolytic myonuclei

Transcriptome of muscle resident cells nuclei was analysed using snRNAseq. Nuclei clusters were identified by specific biomarker expression (Fig. S2A). Nr3c1 (GR) was preferentially expressed in Myh4 (IIb) myonuclei relative to other myonuclei populations (Fig. 2B). Sub-clustering of myonuclei alone resolved fiber-type populations (Myh7, Myh2, Myh1, Myh4) together with NMJ, myotendinous junction, mitochondria-associated nuclei (myonuclei accumulating mt-RNA), and Egr1 positive myonuclei (Fig. 2C). *Ddit4* (REDD1) was also enriched in the Myh4 cluster (IIb) (Fig. 2D). In addition, REDD1 mRNA was also mostly detected by FISH experiment around Myh4-positive myonuclei within Myh4-positive isolated fibers in comparison to Myh4-negative fibers (Fig. 2E). Together, these data show that REDD1 and GR are co-enriched in the fast glycolytic fibers more sensitive to DEX-induced atrophy.

**Figure 2.**
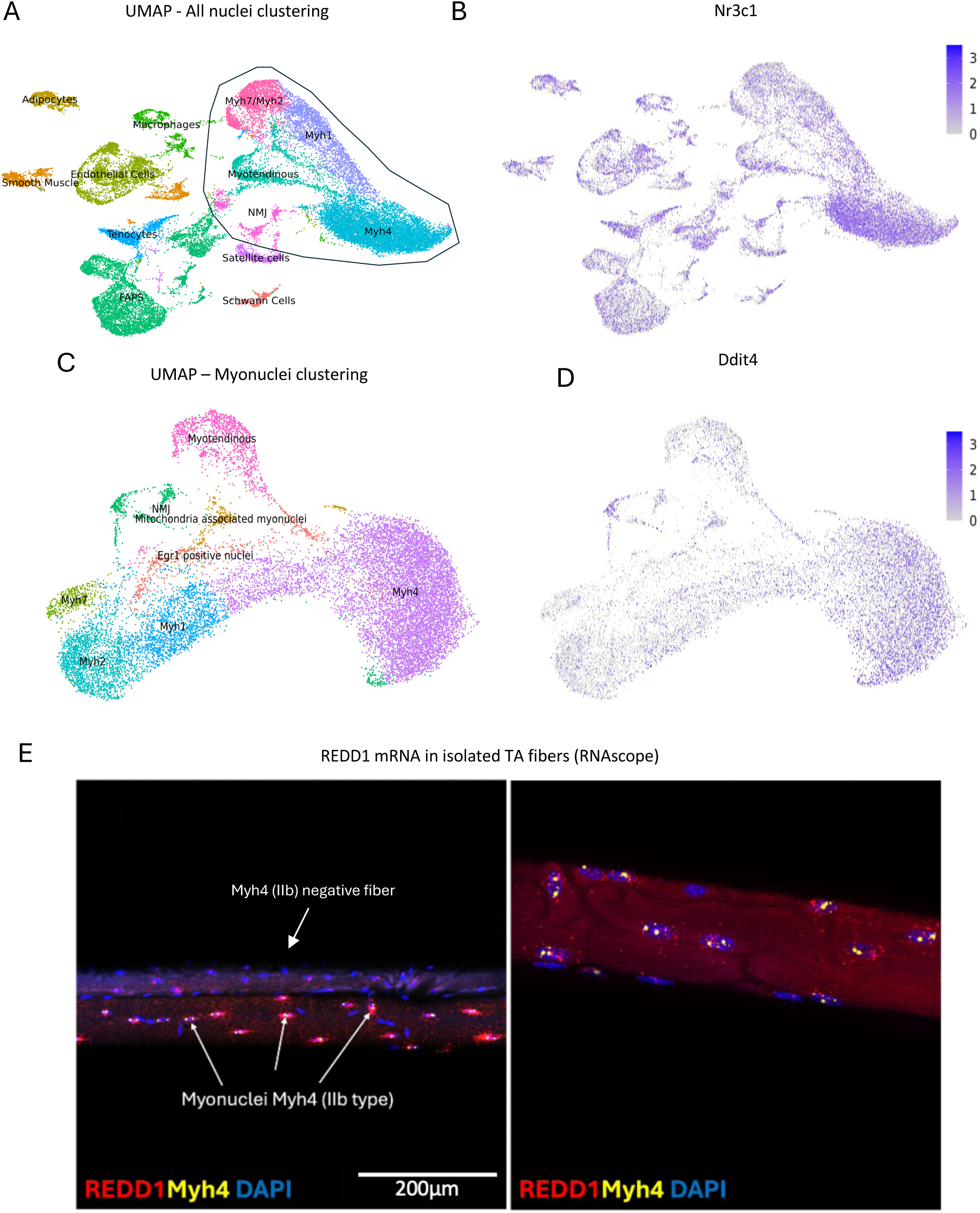
REDD1 (*Ddit4*) transcription is enriched in fast-twitch (Myh4/IIb) myonuclei, which express the glucocorticoid receptor. **A.** UMAP projection of all nuclei from single-nucleus RNA-sequencing of gastrocnemius muscle pooled across the four groups (WT CTL/WT DEX/KO CTL/KO DEX) colored by cluster identity. Non-myogenic populations (adipocytes, macrophages, endothelial cells, smooth muscle, tenocytes, FAPs, satellite cells, Schwann cells) and myonuclear populations (Myh4, Myh1, Myh7/Myh2, myotendinous, NMJ) were resolved. **B.** Feature plot showing the distribution of *Nr3c1* expression across all nuclei; color intensity indicates normalized transcript expression. **C.** UMAP projection of the re-clustered myonuclei, colored by cluster identity: Myh4, Myh1, Myh2, Myh7, myotendinous, NMJ, mitochondria-associated myonuclei, and Egr1-positive nuclei. Only myonuclei were included in this re-clustering analysis. **D.** Feature plot showing the distribution of *Ddit4* expression across myonuclei; color intensity indicates normalized transcript expression. **E.** Representative RNAscope images of isolated TA fibers showing REDD1 (*Ddit4*) mRNA (red), Myh4 pre-mRNA (yellow), and nuclei (DAPI, blue).

### REDD1 deletion in myofiber does not impact Akt/mTOR pathway following 7 days of DEX

To measure protein synthesis, puromycin incorporation was assessed by the SUnSET method. DEX reduced puromycin incorporation by approximately 70% in WT muscle (Fig. 3A). In KO, protein synthesis was already reduced under basal conditions and was not affected by DEX (Fig. 3A). Chronic DEX treatment increased Akt phosphorylation at Thr308 in WT muscle only. mTORC1 targets S6 (Ser235), ULK1 (Ser757) and 4EBP1 (Thr37/46) showed no statistical variation in both WT and KO mice (Fig. 3B). AMPK phosphorylation at Thr172 was increased by DEX in both WT and KO mice muscle (Fig. S3A). AMPK-dependent phosphorylation of ULK1 was reduced only in KO DEX mice (Fig. S3A). Finally, no variations were observed in LC3, p62 or ubiquitination profile (Fig. S3A). These results suggest that in our timing of treatment, protein synthesis inhibition by DEX could not be explained by deregulation of Akt/mTOR pathway.

**Figure 3.**
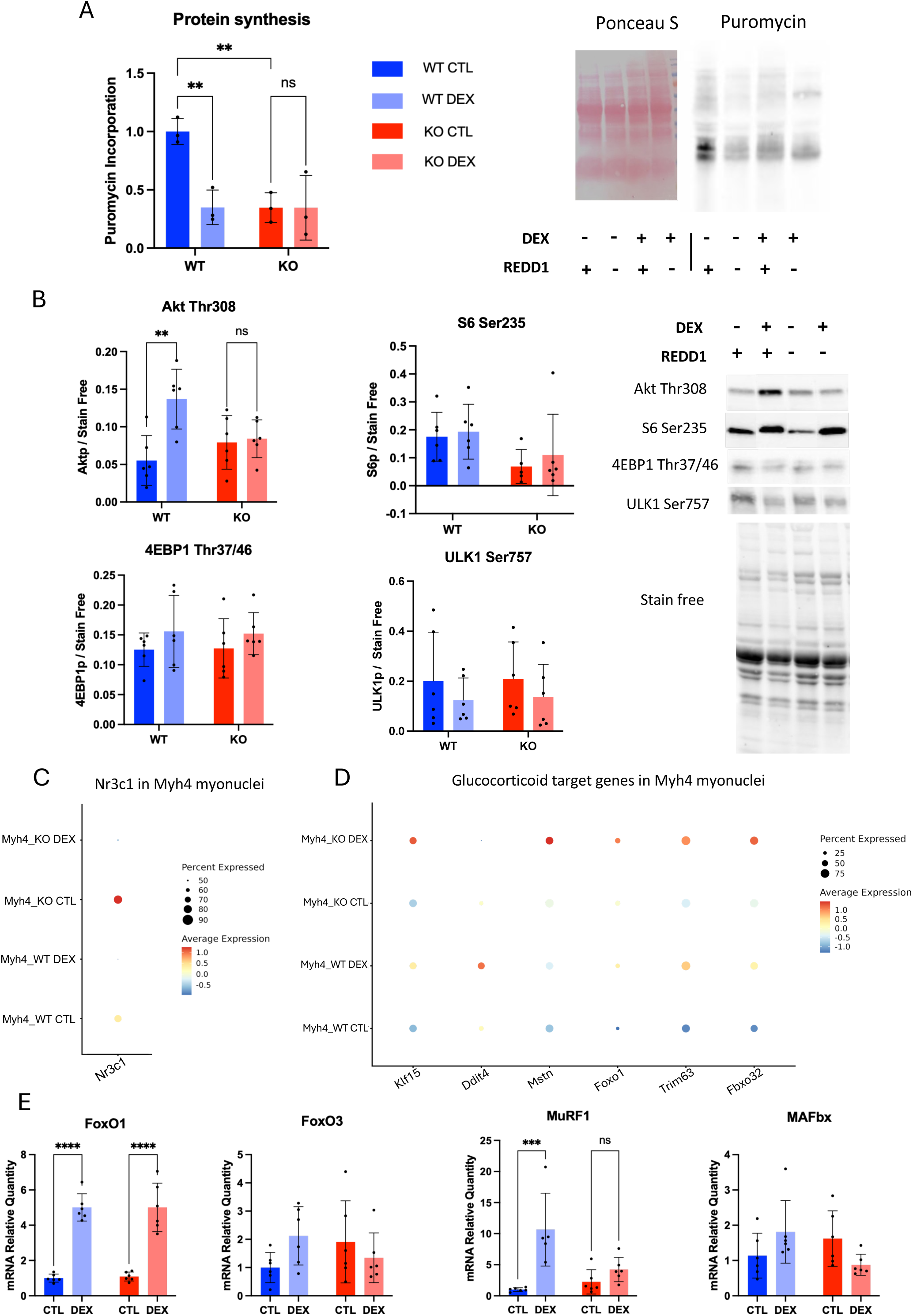
REDD1 deletion uncouples protein synthesis from Akt–mTORC1 signalling and blunts the glucocorticoid transcriptional atrophy program in fast-twitch myonuclei. **A.** Protein synthesis measured by puromycin incorporation (SUnSET) in quadriceps muscle across the four groups, with a representative Ponceau S and anti-puromycin blot. n = 6 per group; two-way ANOVA, Tukey’s post hoc. **B.** Representative western blots and quantification of phosphorylated Akt (Thr308), S6 (Ser235), 4E-BP1 (Thr37/46) and ULK1 (Ser757) in quadriceps of WT and KO mice under CTL and DEX. Phosphorylated signal was normalized to Stain-Free. n = 6 per group; two-way ANOVA, genotype × treatment, Tukey’s post-hoc. **C.** Dot plot of *Nr3c1* (glucocorticoid receptor) expression in Myh4 myonuclei across the four groups (snRNA-seq) Dot size indicates the proportion of expressing nuclei and color intensity indicates normalized expression level. **D.** Dot plot of glucocorticoid target genes (*Klf15*, *Ddit4*, *Mstn*, *Foxo1*, *Trim63*, *Fbxo32*) in Myh4 myonuclei across the four groups (snRNA-seq). Dot size indicates the proportion of expressing nuclei and color intensity indicates normalized expression level. **E.** RT-qPCR of *Foxo1*, *Foxo3*, *Trim63* (MuRF1) and *Fbxo32* (MAFbx) mRNA in quadriceps of WT and KO mice under CTL and DEX. n = 6 per group; two-way ANOVA, Tukey’s post-hoc. mRNA expression was normalized to Rps9.

GR-induced expression of atrogene was assessed by snRNAseq specifically in Myh4 myonuclei. Nr3c1 (GR) expression was higher in KO CTL mice (Fig. 3C,D). Fkbp5, a direct GR target, followed a similar pattern, being higher in KO CTL than in WT CTL at baseline and strongly induced by DEX treatment in both genotypes (Fig. S2B). Accordingly to GR expression atrogenes were over-expressed in KO in comparison to WT Myh4 myonuclei in response to DEX (Fig. 3D). Expression of GR targets involved in proteolysis were then confirmed in whole muscle sample by RT-qPCR. *FoxO1* mRNA was induced by DEX in both WT and KO muscle while MuRF1 (Trim63) expression was induced by DEX in WT but not in KO. *FoxO3 and MAFbx* (*Fbxo32*) expressions were not significantly changed in either genotype by DEX (Fig. 3E).

### Compartment-specific remodelling of mitochondria and ER-mitochondria contacts sites under dexamethasone

In WT muscle, dexamethasone reduced mitochondrial area in both the perinuclear (∼0.65-fold) and IMF pools (∼0.67-fold)(Fig. 4A and B). In KO muscle, mitochondrial area was already lower than in WT in both compartments (∼0.5 fold). Unexpectedly, DEX increased mitochondrial area in KO muscle (∼1.8-fold perinuclear; ∼1.6-fold IMF; Fig. 4B). Mitochondrial shape, assessed by eccentricity, was unchanged in the perinuclear pool but increased under DEX in the IMF pool of both genotypes (Fig. 4B). The subsarcolemmal pool, analysed separately, showed no significant change in area or eccentricity (Fig. S3A, B). Additional shape descriptors were quantified in all three compartments: mitochondrial perimeter followed a consistent pattern, being lower in KO CTL than in WT CTL and higher in KO DEX than in KO CTL, while DEX reduced solidity in the perinuclear and IMF pools of WT muscle only (Fig.S3 C, D).

**Figure 4.**
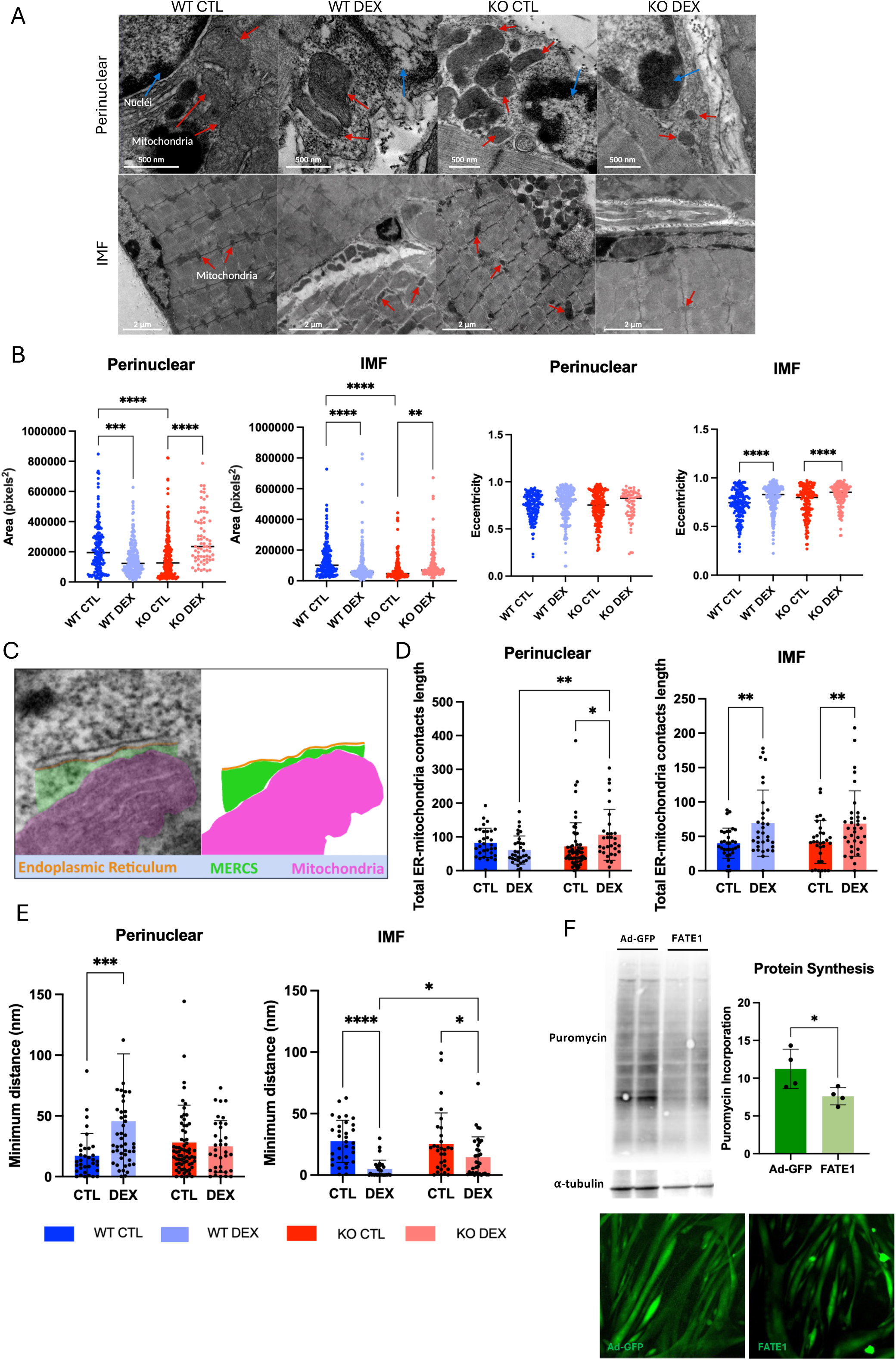
Dexamethasone remodels mitochondrial morphology and ER–mitochondria contacts in a compartment- and genotype-specific manner, and ER–mitochondria uncoupling limits protein synthesis in vitro. **A.** Representative transmission electron micrographs of the perinuclear and intermyofibrillar (IMF) compartments in TA muscle of WT and KO mice under CTL and DEX. Mitochondria are indicated by red arrows and nuclei by blue arrows. **B.** Quantification of mitochondrial area in pixels² and eccentricity in the perinuclear and IMF compartments across the four groups. n = 3 to 5 mice per group; between ∼70 and ∼220 mitochondria analysed per group; two-way ANOVA, Tukey’s post-hoc. **C.** Representative electron micrograph of a mitochondria-associated endoplasmic reticulum membrane (MERCS, in green), showing apposed ER (orange) and mitochondrion (pink). **D.** Total ER–mitochondria contact length in the perinuclear and IMF compartments, WT and KO, under CTL and DEX. Two-way ANOVA, Tukey’s post-hoc. n= 3 to 5 mice per group; between ∼∼30 and ∼60 MERCS quantified per group. **E.** Minimum ER–mitochondria distance (nm) in the perinuclear and IMF compartments, WT and KO, under CTL and DEX. n= 3 to 5 mice per group; between ∼30 to ∼60 MERCS quantified per group. Two-way ANOVA, Tukey’s post-hoc. **F.** C2C12 myotubes transduced with control adenovirus (Ad-GFP) or FATE1-expressing adenovirus. Left: representative anti-puromycin and α-tubulin western blot. n = 3 per group; unpaired t-test. Bottom: representative GFP fluorescence of Ad-GFP and FATE1 myotubes confirming transduction.

We next quantified MERCS, defined by regions where mitochondria and ER are close (Fig. 4C). The total length of ER-mitochondria contacts increased under dexamethasone treatment in the IMF pool of both genotypes (Fig. 4D). In the perinuclear pool, this increase was significant only in KO muscle (Fig. 4E). In the IMF pool, DEX reduced minimum distance between organelles in both genotypes (from ∼28 to ∼4.8 nm in WT, and from ∼25 to ∼14.5nm in KO), although this reduction tended to be smaller in KO than in WT group (p interaction=0.058). In the perinuclear pool, DEX increased the minimum distance in WT (from ∼18 to ∼45 nm) (Fig. 4E). The full distribution of contact lengths across distance is provided in Supplementary Fig. 3E. Together, these data indicate that REDD1 is important for DEX-induced remodelling of MERCS and mitochondrial network in a subcellular compartment-specific manner.

To test whether the integrity of MERCS influences protein synthesis, C2C12 myotubes were infected with an adenovirus expressing Ad-FATE1-GFP, which uncouples ER-mitochondria contacts, or with Ad-GFP control adenovirus (Fig. 4F). FATE1 expression reduced puromycin incorporation relative to control, indicating that disrupting MERCS is sufficient to lower protein synthesis in muscle cells.

In WT mice only, in addition to MERCS alteration, DEX increased distance between mitochondria and nuclei (Fig. 5A and B). This alteration of mitochondrial contacts with nuclei by DEX was associated with a reduction of the number of nuclei accumulating mt-RNA (KO CTL = 28,9%, KO DEX = 34,3%, WT CTL = 25%, WT DEX = 11,8%) (Fig. S5A, B, C). Gene expression comparison and ingenuity pathway analysis of this specific cluster showed that these nuclei were enriched for a set of sarcomeric, cytoskeletal and extracellular matrix transcripts (Fig. S5 D, E). These results suggest that REDD1 regulate mitochondrial association with nuclei that may influence anabolic transcription program.

**Figure 5.**
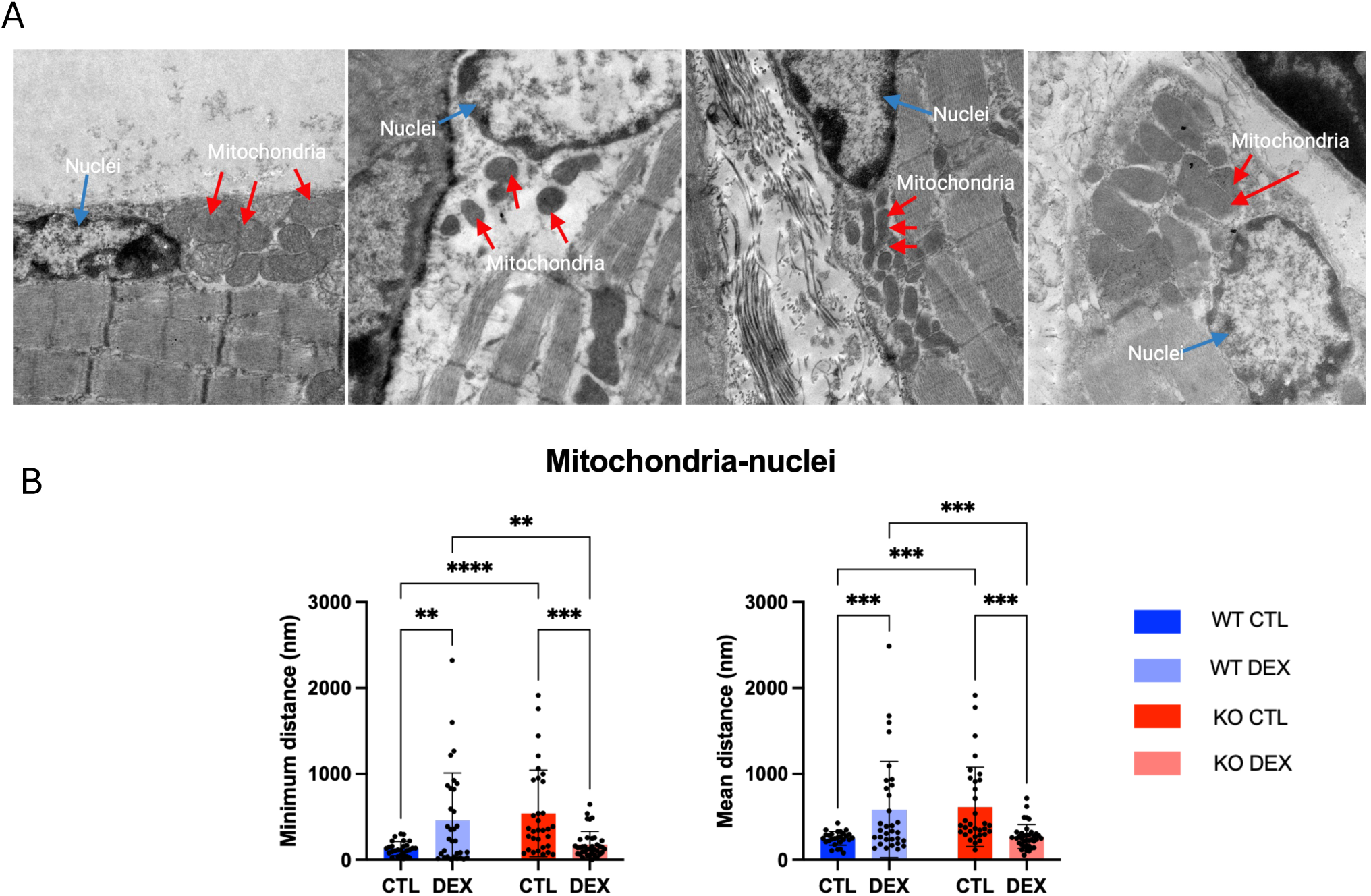
REDD1 deletion protects from DEX-induced increase of mitochondrial nuclei distance\. **A.** Representative transmission electron microscopy (TEM) images of the perinuclear region in WT and KO muscle under CTL and DEX. Blue labels indicate the nucleus ; red arrows point to mitochondria in close apposition to the nuclear envelope (perinuclear mitochondria– ER/nuclear contact sites). **D.** Quantification of the minimum distance (left) and mean distance (right, nm) between mitochondria and the nuclear envelope in WT and KO muscle under CTL and DEX. Two-way ANOVA. Tukey’s post hoc.

### REDD1 localizes to the NMJ, where MERCS control acetylcholine receptor clustering

DEX reduced the endplate area in WT mice only (Fig. 6A). In KO muscle, the endplate area was already lower than in WT at baseline and did not decrease with DEX (Fig. 6A). In addition, DEX treated WT mice displayed higher NMJ fragmentation in comparison to DEX treated KO mice (Fig. S6 A).

**Figure 6.**
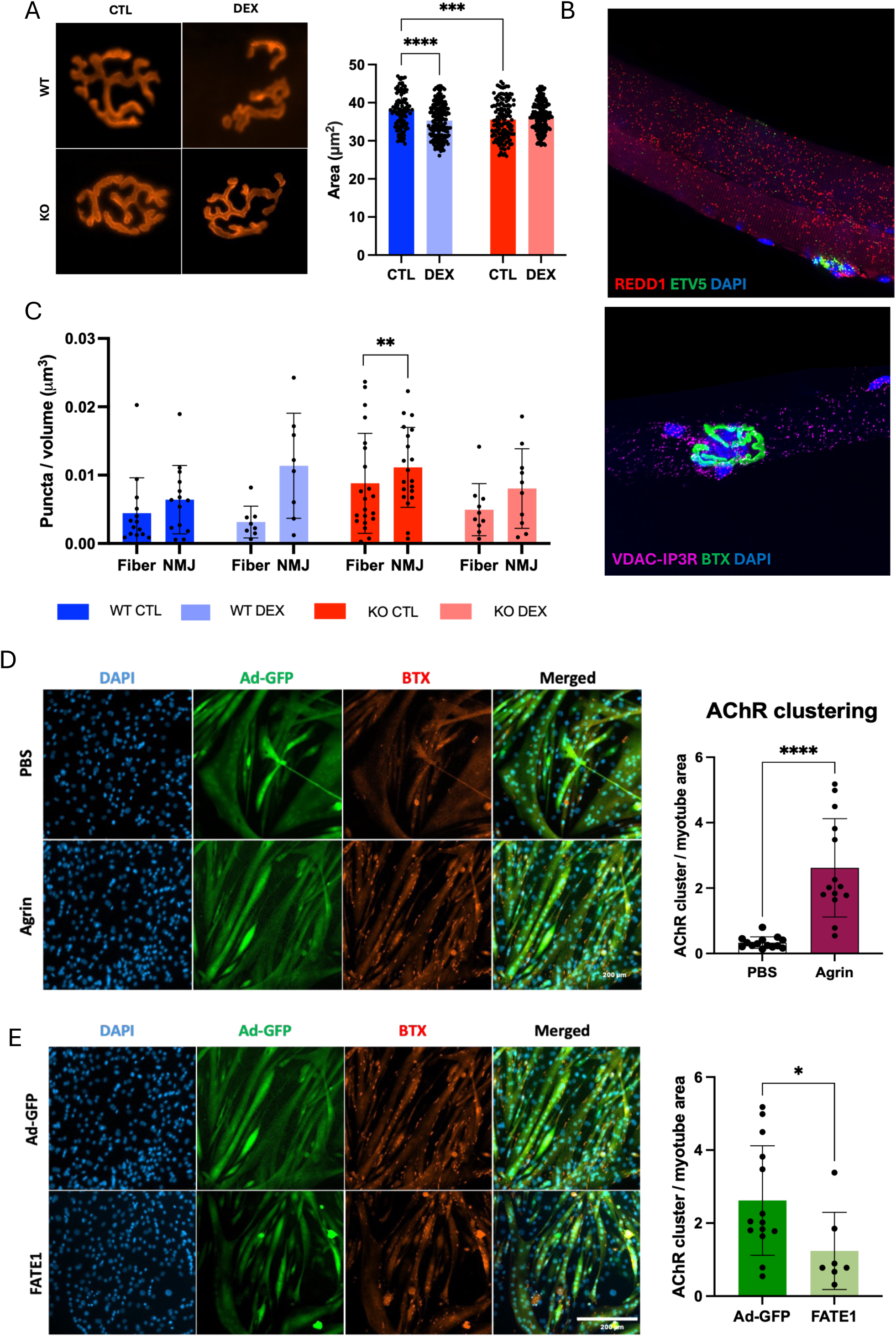
REDD1 is enriched at the neuromuscular junction, while experimentally increasing ER–mitochondria distance reduces acetylcholine receptor clustering in vitro. **A.** Representative images of neuromuscular junctions (NMJs) stained with α-bungarotoxin (BTX) in WT and KO mice under CTL and DEX, with quantification of NMJ area (µm²). n = 120 to 200 NMJs from 6 mice per group; two-way ANOVA, Tukey’s post-hoc. **B.** Representative RNAscope image of an isolated TA fiber showing REDD1 (*Ddit4*) mRNA (red), the synaptic marker ETV5 (green), and nuclei (DAPI, blue). **C.** Quantification of ER–mitochondria contact density (VDAC–IP3R PLA puncta per µm³) at synaptic fiber regions versus the NMJ, in WT and KO mice under CTL and DEX. Data were analyzed by three-way mixed-effects analysis (REML) with location (fiber *vs* NMJ) as a repeated within-subject factor, treatment and genotype as between-group factors. Main effect of location p = 0.0003. Right: proximity ligation assay (PLA) signal (magenta), BTX (green), DAPI (blue). **D.** C2C12 myotubes treated with PBS or agrin, stained for DAPI (blue), Ad-GFP (green) and BTX (red; AChR), with merged panels. Right: quantification of AChR clusters per myotube area. Unpaired t-test, ****p<0.0001. **E.** C2C12 myotubes transduced with Ad-GFP or FATE1 adenovirus, same staining. Right: quantification of AChR clusters per myotube area. Unpaired t-test *p < 0.05. AChR clusters were quantified as BTX-positive clusters normalized to myotube area.

FISH and snRNAseq analysis highlighted that REDD1 and GR are expressed in NMJ (Fig. 6B and Fig. S6B). Atrogenes were induced by DEX in NMJ in both genotypes (Fig. S6C). Ingenuity pathway analysis of NMJ myonuclei transcriptome indicated that, under DEX, acetylcholine receptor and agrin signalling were more activated in KO than in WT suggesting that REDD1 is necessary DEX-induced inhibition of NMJ transcriptomic program (Fig. S6D).

MERCS density, measured by PLA targeting IP3R and VDAC, was higher at the NMJ than in fiber body (main effect of location, p=0.0003)(Fig. 6C). This identifies the NMJ as a compartment enriched in MERCS. To test whether MERCS are involved in AChR clustering, we used C2C12 myotubes submitted to an agrin treatment to induce AChR clustering and quantified as BTX-positive clusters per myotube area (Fig. 6D). Disruption of MERCS by Ad-FATE1 decreased agrin-induced AChR clustering relative to the Ad-GFP control (Fig. 6E).

## Discussion

We studied the contribution of REDD1 expression in skeletal muscle on glucocorticoid-induced fast glycolytic fiber atrophy using a tamoxifen-inducible, muscle specific knockout. Our main findings are that REDD1 promotes glycolytic fibers atrophy and NMJ destabilization in response to glucocorticoids treatment. This REDD1 dependent atrophy is associated with alteration of mitochondrial morphology and association with other organelles such as endoplasmic reticulum and nuclei as well as modification of transcriptome and protein synthesis. We notably show for the first time that MERCS integrity is necessary to maintain protein synthesis and to promote acetylcholine receptors aggregation. Finally, our results also demonstrate that REDD1 is necessary to maintain normal mitochondrial morphology and protein synthesis at basal state.

### Loss of REDD1 induces basal muscle atrophy

REDD1-deficient muscle displayed a lower baseline quadriceps mass, lower Myh4 fiber CSA, basal mitochondrial/MERCS alteration and lower baseline rate of protein synthesis than WT. One hypothesis is that this baseline remodelling of mitochondrial network reduces mitochondrial efficiency and thereby limits the energy available for an ATP-costly processes such as protein synthesis. In fact, a myriad of studies showed that mitochondria morphology and MERCS alteration impact mitochondrial function (30–32) as well as protein synthesis (12). These results are in contradiction with our previous studies performed in REDD1 total KO mice (19,21). Indeed, in contrast to the muscle-specific and inducible REDD1 KO mice, muscle mitochondrial function, muscle protein synthesis and basal muscle mass are not altered in young total KO mice (19,21). In the whole body and constitutive KO model, muscle atrophy occurs only in old animals (12 months old) while muscle wasting is observable only 3 weeks after REDD1 deletion in inducible and muscle specific KO model (21). These results suggest that compensatory mechanisms exist in the total KO to compensate the absence of REDD1.

### REDD1 and GR mark the fast fiber atrophy

DEX preferentially reduced the cross-sectional area of the most glycolytic fibers IIb while IIa fibers are less affected. This reproduces the unexplained fiber selectivity of glucocorticoid myopathy (1,2). Our snRNAseq and FISH data showed that both REDD1 and GR were enriched in IIb myonuclei, suggesting that GR/REDD1 pathway is involved in their atrophy. In fact, REDD1 is a direct target of GR and has been shown to promote global muscle atrophy in response to DEX treatment (18,19). Confirming this hypothesis, our results showed that REDD1 deletion in fibers spared type IIb fibers from DEX-induced atrophy.

Glucocorticoid treatment are known to affect protein balance (33,34). In our conditions, protein synthesis is reduced by ∼70% in WT muscle, yet the master regulator of protein synthesis, Akt/mTORC1 pathway (S6 Ser235, 4EBP1 Thr37/46, Ulk1 Ser757), did not follow this fall. In muscle specific REDD1 KO muscle, synthesis was already low at baseline without alteration of mTOR pathway, and we observed no additional change in response to DEX. Interestingly, we have previously demonstrated that REDD1 regulates protein synthesis in response to acute DEX treatment through regulation of Akt/mTORC1 pathway (19). The results of this present study on chronic DEX treatment suggest that REDD1-dependent regulation of protein synthesis could occur through other mechanisms. Decorrelation between REDD1/protein synthesis and mTORC1 activation has already been observed such as during overload-induced hypertrophy, hypoxia or fasting (9,42). REDD1-dependent inhibition of protein synthesis could reflect a bioenergetic limit, insufficient ATP to sustain a high rate of translation, which would link it to the mitochondrial and energetic phenotype described below. In agreement with this hypothesis, mitochondrial function alteration drives to muscle atrophy (36). Moreover, we have shown in this study that alteration of MERCS is associated with reduced protein synthesis.

### Compartment-specific remodelling of Mitochondria and ER-mitochondria Contacts Sites (MERCS)

Skeletal muscle mitochondria present different pools that differ by their localization in the cell: intermyofibrillar (IMF), perinuclear and subsarcolemmal pool. Recent studies suggest that classical mitochondrial classification does not fully capture the spatial and functional heterogeneity of skeletal muscle mitochondria, highlighting the potential value of refining mitochondrial subpopulations based on their precise subcellular localization and proximity to other cellular structures such as vessels or nuclei (37). The same heterogeneity should apply to their ER contacts, which already differ in frequency and extent between pools: different mitochondrial populations likely form different MERCS, with different local functions(37,38).

In WT mice, DEX treatment reduced mitochondrial area in both pools. Mitochondrial size reduction is compatible with the mitochondrial fragmentation induced by glucocorticoids and occurring before atrogene program activation (10,11). In REDD1 KO muscle, mitochondria were smaller at baseline in both compartments and unexpectedly DEX treatment increased their area. REDD1 is thus required for mitochondrial fragmentation under DEX treatment, that is consistent with the role of REDD1 in autophagy and mitochondrial quality control (39).

Because REDD1 is expressed in MERCS subcellular fraction and engages with VDAC-IP3R-GRP75 tethering complexes (21), MERCS were also analysed in a compartment specific manner. Interpreting distance changes requires understanding how spacing affects function of MERCS. TEM places physiological tethers at ∼10 nm at the smooth ER and ∼25nm at the rough ER (9). This spacing is important for Ca^2+^ handling: synthetic linkers fixing distance tethering show that IP3R mediated transfer is optimal near 20 nm and declines when the interface becomes either too large (30 nm) or too narrow (10 nm) inhibiting Ca^2+^ flux (40). Beyond Ca^2+^, enhanced ER-mitochondria tethering may influence cellular protein homeostasis (12). In this context, the IMF-ER minimum distance was measured near the functional range at baseline in both WT and KO mice (WT CTL ∼27.5nm: KO CTL 25.2nm). DEX shortened distance in both genotypes, reaching ∼4.8nm in WT and ∼14.5 nm in KO. Although both genotypes tightened, only in WT DEX muscles did the gap fall below 10 nm, into the range where Ca^2+^ transfer becomes inefficient and the contacts are likely too close to sustain the normal metabolic exchange. That indicates that REDD1 deletion could protect IMF function in response to DEX. In fact, we have previously shown that REDD1 total KO mice are resistant to DEX-induced alteration of mitochondrial respiration (21).

The perinuclear pool of mitochondria has both bioenergetic and translational relevance as in skeletal muscle ribosomes and mRNA concentrate around myonuclei (41,42). In contrast to the IMF pool, DEX treatment increased the mitochondria-ER distance on the perinuclear compartment of WT muscle only. Because MERCS regulate ATP supply (11,43), REDD1 dependent change in perinuclear contact could alter local energy available for perinuclear translation. These results are in line with previous studies showing that REDD1 silencing in vitro leads to increase of MERCS extent, ATP content and protein synthesis (21,44). Moreover, REDD1 expression during exercise is associated with concomitant reduction of MERCS and protein synthesis (45). Proving the role of MERCS in protein synthesis, we showed that MERCS spacing by overexpression of FATE1 was sufficient to reduce protein synthesis.

In addition, DEX also increased the distance between mitochondria and nuclei that is associated with a reduction of the number of myonuclei enriched for mt-RNA displaying specific anabolic transcriptome only in WT mice. These nuclei overexpress sarcomeric, cytoskeletal and extracellular matrix genes in comparison to other myonuclei. This result suggests that REDD1 could be important to regulate translation potentially by controlling mitochondrial energetics near nuclei. In agreement with this proposition, recent study shows that mitochondrial association with nuclear pore sustains nuclear ATP levels, supports transcription, histone modification and chromatin dynamics (46). In addition, other studies also indicate that mt-RNA could play a role in the regulation of gene expression by fixation with gene promotor (47). Therefore, REDD1 may control transcriptome by regulating ATP supply to nuclei and potentially by modifying mt-RNA present in nuclei. This hypothesis is supported by the strong modification of myonuclei transcription observed in our snRNAseq data of Ko mice (not shown). These results on transcription open new questions about the role of REDD1 and MERCS in the regulation of nuclei energetic and transcriptome and maybe translatome. Further studies are needed to clarify this point.

### REDD1 and MERCS at the neuromuscular junction

We showed here that REDD1 is expressed in NMJ and that its deletion protected NMJ from DEX treatment-induced reduction of area and fragmentation. The NMJ concentrates mitochondria and myonuclei and sustains high local synthesis of continually renewed synaptic protein such as AChR subunits (16,48,49). Consistent with that demand, VDAC-IP3R contacts were denser at the NMJ than along the fiber suggesting an important role of MERCS in NMJ. Disrupting MERCS with FATE1 reduced agrin-induced AChR clustering in C2C12 cells, providing a causal link between MERCS integrity and postsynaptic AChR assembly. Pathway analysis of the NMJ myonuclei indicated that, under DEX treatment, AChR signalling was more active in KO than WT, which may reflect a compensatory postsynaptic response. Taken together, these results suggest that REDD1 is required for DEX-induced NMJ alteration potentially through disruption of MERCS and NMJ transcriptomic program.

## Conclusion

REDD1 is required for DEX induced fast fiber atrophy and NMJ destabilization acting on mitochondrial network (depending of subcellular domain) and synthesis process including transcription and translation. Together, our data raise the question of the role of REDD1-dependent remodelling of the perinuclear mitochondrial network in the mechanism of glucocorticoid-induced muscle wasting. In addition, our study also show that REDD1 is important for the control of mitochondrial network and protein synthesis at basal state. Fast-fiber atrophy is a shared feature of cancer cachexia, obesity, SLA, where it contributes to loss of mobility and worse outcomes (50–52). Importantly, these pathologies are also associated with REDD1 expression suggesting that the mechanisms described in our study could maybe be extended to other pathologies affecting fast fibers and NMJ opening therapeutic strategies perspectives (50,53–55).

## Acknowledgement

We acknowledge GENOM’IC, PIME and MOUSET’IC core facilities, and the animal core facility of Institut Cochin. We thank the animal staff from RAM-Metamus DMEM facility for technical support and animal care expertise (https://dmem.montpellier.hub.inrae.fr/).

## Language assistance

Claude AI was used marginally to refine English in selected passages and to help formulate concise, impact-full text in sections with strict word limits.

## References

1. Dekhuijzen PN, Gayan-Ramirez G, Bisschop A, De Bock V, Dom R, Decramer M. Corticosteroid treatment and nutritional deprivation cause a different pattern of atrophy in rat diaphragm. J Appl Physiol. 1 févr 1995;78(2):629–37. doi:10.1152/jappl.1995.78.2.629

2. Schakman O, Gilson H, Thissen JP. Mechanisms of glucocorticoid-induced myopathy. J Endocrinol. avr 2008;197(1):1–10. doi:10.1677/JOE-07-0606

3. Schakman O, Kalista S, Barbé C, Loumaye A, Thissen JP. Glucocorticoid-induced skeletal muscle atrophy. Int J Biochem Cell Biol. oct 2013;45(10):2163–72. doi:10.1016/j.biocel.2013.05.036

4. Liu J, Peng Y, Wang X, Fan Y, Qin C, Shi L, et al. Mitochondrial Dysfunction Launches Dexamethasone-Induced Skeletal Muscle Atrophy via AMPK/FOXO3 Signaling. Mol Pharm. 4 janv 2016;13(1):73–84. doi:10.1021/acs.molpharmaceut.5b00516

5. Troncoso R, Paredes F, Parra V, Gatica D, Vásquez-Trincado C, Quiroga C, et al. Dexamethasone-induced autophagy mediates muscle atrophy through mitochondrial clearance. Cell Cycle. 15 juill 2014;13(14):2281–95. doi:10.4161/cc.29272

6. Dong H, Tsai SY. Mitochondrial PrOPERTIES in Skeletal Muscle Fiber. Cells. 30 août 2023;12(17):2183. doi:10.3390/cells12172183

7. Buttgereit F, Brand MD. A hierarchy of ATP-consuming processes in mammalian cells. Biochem J. 15 nov 1995;312(1):163–7. doi:10.1042/bj3120163

8. Lynch M, Marinov GK. The bioenergetic costs of a gene. Proc Natl Acad Sci. 22 déc 2015;112(51):15690–5. doi:10.1073/pnas.1514974112

9. Csordás G, Renken C, Várnai P, Walter L, Weaver D, Buttle KF, et al. Structural and functional features and significance of the physical linkage between ER and mitochondria. J Cell Biol. 25 sept 2006;174(7):915–21. doi:10.1083/jcb.200604016

10. Denton RM. Regulation of mitochondrial dehydrogenases by calcium ions. Biochim Biophys Acta BBA - Bioenerg. nov 2009;1787(11):1309–16. doi:10.1016/j.bbabio.2009.01.005

11. Szabadkai G, Bianchi K, Várnai P, De Stefani D, Wieckowski MR, Cavagna D, et al. Chaperone-mediated coupling of endoplasmic reticulum and mitochondrial Ca2+ channels. J Cell Biol. 18 déc 2006;175(6):901–11. doi:10.1083/jcb.200608073

12. Tonelli E, Malecka J, Barberis E, Romano C, Pessolano E, Talmon M, et al. Remodelling of Cellular Protein Homeostasis by Enhanced ER-Mitochondrial Tethering. Contact J Inter-Organelle Commun. avr 2025;8:25152564251329704. doi:10.1177/25152564251329704

13. Fahim MA. Chronic corticosterone treatment-induced ultrastructural changes at rat neuromuscular junction. Anat Rec. juill 1995;242(3):424–31. doi:10.1002/ar.1092420315

14. Tomas I Ferre J, Mayayo E, Fenoll I Brunet R. Morphometric study of the neuromuscular synapses in the adult rat with special reference to the remodelling concept. Biol Cell. janv 1987;60(2):133–44. doi:10.1111/j.1768-322X.1987.tb00553.x

15. Bruneau EG, Akaaboune M. The dynamics of recycled acetylcholine receptors at the neuromuscular junction in vivo. Development. 15 nov 2006;133(22):4485–93. doi:10.1242/dev.02619

16. Altman T, Geller D, Kleeblatt E, Gradus-Perry T, Perlson E. An *in vitro* compartmental system underlines the contribution of mitochondrial immobility to the ATP supply in the NMJ. J Cell Sci. 1 déc 2019;132(23). doi:10.1242/jcs.234492

17. Guzman SD, Fraczek PM, Itsani K, Furati EK, Juros D, Kenney G, et al. Age-Associated Dysregulation of Postsynaptic Mitochondria Perturbs Reinnervation Kinetics. Aging Cell. janv 2026;25(1):e70355. doi:10.1111/acel.70355

18. Wang H, Kubica N, Ellisen LW, Jefferson LS, Kimball SR. Dexamethasone Represses Signaling through the Mammalian Target of Rapamycin in Muscle Cells by Enhancing Expression of REDD1. J Biol Chem. déc 2006;281(51):39128–34. doi:10.1074/jbc.m610023200

19. Britto FA, Begue G, Rossano B, Docquier A, Vernus B, Sar C, et al. REDD1 deletion prevents dexamethasone-induced skeletal muscle atrophy. Am J Physiol-Endocrinol Metab. 1 déc 2014;307(11):E983–93. doi:10.1152/ajpendo.00234.2014

20. Gordon BS, Williamson DL, Lang CH, Jefferson LS, Kimball SR. Nutrient-induced stimulation of protein synthesis in mouse skeletal muscle is limited by the mTORC1 repressor REDD1. J Nutr. avr 2015;145(4):708–13. doi:10.3945/jn.114.207621 PubMed PMID: 25716553; PubMed Central PMCID: PMC4381770.

21. Britto FA, Cortade F, Belloum Y, Blaquière M, Gallot YS, Docquier A, et al. Glucocorticoid-dependent REDD1 expression reduces muscle metabolism to enable adaptation under energetic stress. BMC Biol. déc 2018;16(1):65. doi:10.1186/s12915-018-0525-4

22. Notini AJ, McClive PJ, Meachem SJ, Van Den Bergen JA, Western PS, Gustin SE, et al. Redd1 Is a Novel Marker of Testis Development but Is Not Required for Normal Male Reproduction. Sex Dev. 2012;6(5):223–30. doi:10.1159/000339723

23. Di Gallo M, Guilbert T, Pereira D, Cepella Z, Braud-Mussi R, Jauliac E, et al. FLASH: innovative integrated enzymatic-fluorescent labeling for automated muscle fiber typing, metabolic and morphometric analysis. Skelet Muscle. 21 nov 2025;15(1):32. doi:10.1186/s13395-025-00401-6

24. Minty G, Hoppen A, Boehm I, Alhindi A, Gibb L, Potter E, et al. aNMJ-morph: a simple macro for rapid analysis of neuromuscular junction morphology. R Soc Open Sci. avr 2020;7(4):200128. doi:10.1098/rsos.200128

25. Theurey P, Tubbs E, Vial G, Jacquemetton J, Bendridi N, Chauvin MA, et al. Mitochondria-associated endoplasmic reticulum membranes allow adaptation of mitochondrial metabolism to glucose availability in the liver. J Mol Cell Biol. avr 2016;8(2):129–43. doi:10.1093/jmcb/mjw004

26. Hucteau E, Hureau TJ, Pagano AF, Mallard J. AMITO: An open-access application for assisted analysis of mitochondria in transmission electron microscopy images. Am J Physiol-Cell Physiol. 18 août 2026;ajpcell.00300.2026. doi:10.1152/ajpcell.00300.2026

27. Macaux G, Di Gallo M, Taglietti V, Amthor H, Maire P. Cell-Hub: a graphical interface for end-to-end single-cell RNA sequencing analysis [Internet]. Bioinformatics; 2026 [cité 24 août 2026]. Disponible sur: http://biorxiv.org/lookup/doi/10.64898/2026.07.15.738621 doi:10.64898/2026.07.15.738621

28. Owen DB, Chamberlain KT, Shishido S, Grow WA. Ethanol decreases agrin-induced acetylcholine receptor clustering in C2C12 myotube culture. Toxicol In Vitro. mars 2010;24(2):645–51. doi:10.1016/j.tiv.2009.09.020

29. Chang YF, Chou HJ, Yen YC, Chang HW, Hong YR, Huang HW, et al. Agrin induces association of *Chrna1* mRNA and nicotinic acetylcholine receptor in C2C12 myotubes. FEBS Lett. 21 sept 2012;586(19):3111–6. doi:10.1016/j.febslet.2012.07.068

30. Cárdenas C, Miller RA, Smith I, Bui T, Molgó J, Müller M, et al. Essential Regulation of Cell Bioenergetics by Constitutive InsP3 Receptor Ca2+ Transfer to Mitochondria. Cell. juill 2010;142(2):270–83. doi:10.1016/j.cell.2010.06.007

31. Bravo R, Vicencio JM, Parra V, Troncoso R, Munoz JP, Bui M, et al. Increased ER– mitochondrial coupling promotes mitochondrial respiration and bioenergetics during early phases of ER stress. J Cell Sci. 1 juill 2011;124(13):2143–52. doi:10.1242/jcs.080762

32. Marchi S, Patergnani S, Pinton P. The endoplasmic reticulum–mitochondria connection: One touch, multiple functions. Biochim Biophys Acta BBA -Bioenerg. avr 2014;1837(4):461–9. doi:10.1016/j.bbabio.2013.10.015

33. Löfberg E, Gutierrez A, Wernerman J, Anderstam B, Mitch WE, Price SR, et al. Effects of high doses of glucocorticoids on free amino acids, ribosomes and protein turnover in human muscle. Eur J Clin Invest. mai 2002;32(5):345–53. doi:10.1046/j.1365-2362.2002.00993.x

34. Tomas FM, Munro HN, Young VR. Effect of glucocorticoid administration on the rate of muscle protein breakdown *in vivo* in rats, as measured by urinary excretion of *N* τ-methylhistidine. Biochem J. 15 janv 1979;178(1):139–46. doi:10.1042/bj1780139

35. Gordon BS, Liu C, Steiner JL, Nader GA, Jefferson LS, Kimball SR. Loss of REDD1 augments the rate of the overload-induced increase in muscle mass. Am J Physiol-Regul Integr Comp Physiol. 1 sept 2016;311(3):R545–57. doi:10.1152/ajpregu.00159.2016

36. Hyatt HW, Powers SK. Mitochondrial Dysfunction Is a Common Denominator Linking Skeletal Muscle Wasting Due to Disease, Aging, and Prolonged Inactivity. Antioxidants. 11 avr 2021;10(4):588. doi:10.3390/antiox10040588 PubMed PMID: 33920468; PubMed Central PMCID: PMC8070615.

37. Parry HA, Willingham TB, Giordano KA, Kim Y, Qazi S, Knutson JR, et al. Impact of capillary and sarcolemmal proximity on mitochondrial structure and energetic function in skeletal muscle. J Physiol. mai 2024;602(9):1967–86. doi:10.1113/JP286246

38. Boncompagni S, Rossi AE, Micaroni M, Beznoussenko GV, Polishchuk RS, Dirksen RT, et al. Mitochondria Are Linked to Calcium Stores in Striated Muscle by Developmentally Regulated Tethering Structures. Parton RG, éditeur. Mol Biol Cell. févr 2009;20(3):1058–67. doi:10.1091/mbc.e08-07-0783

39. Qiao S, Dennis M, Song X, Vadysirisack DD, Salunke D, Nash Z, et al. A REDD1/TXNIP pro-oxidant complex regulates ATG4B activity to control stress-induced autophagy and sustain exercise capacity. Nat Commun. 28 avr 2015;6(1):7014. doi:10.1038/ncomms8014

40. Dematteis G, Tapella L, Casali C, Talmon M, Tonelli E, Reano S, et al. ER-mitochondria distance is a critical parameter for efficient mitochondrial Ca2+ uptake and oxidative metabolism. Commun Biol. 10 oct 2024;7(1):1294. doi:10.1038/s42003-024-06933-9

41. Nevalainen M, Kaakinen M, Metsikkö K. Distribution of mRNA transcripts and translation activity in skeletal myofibers. Cell Tissue Res. sept 2013;353(3):539–48. doi:10.1007/s00441-013-1659-x PubMed PMID: 23736382.

42. Pinheiro H, Pimentel MR, Sequeira C, Oliveira LM, Pezzarossa A, Roman W, et al. mRNA distribution in skeletal muscle is associated with mRNA size. J Cell Sci. 15 juill 2021;134(14):jcs256388. doi:10.1242/jcs.256388

43. Betz C, Stracka D, Prescianotto-Baschong C, Frieden M, Demaurex N, Hall MN. mTOR complex 2-Akt signaling at mitochondria-associated endoplasmic reticulum membranes (MAM) regulates mitochondrial physiology. Proc Natl Acad Sci. 30 juill 2013;110(31):12526–34. doi:10.1073/pnas.1302455110

44. Torres-Velarde JM, Kolora SRR, Khudyakov JI, Crocker DE, Sudmant PH, Vázquez-Medina JP. Elephant seal muscle cells adapt to sustained glucocorticoid exposure by shifting their metabolic phenotype. Am J Physiol-Regul Integr Comp Physiol. 1 sept 2021;321(3):R413–28. doi:10.1152/ajpregu.00052.2021

45. Merle A, Jollet M, Britto FA, Goustard B, Bendridi N, Rieusset J, et al. Endurance exercise decreases protein synthesis and ER-mitochondria contacts in mouse skeletal muscle. J Appl Physiol. 1 nov 2019;127(5):1297–306. doi:10.1152/japplphysiol.00196.2019

46. Menendez-Montes I, Marin-Vicente C, Mukherjee S, Ahmed MS, Gomez MJ, Anene-Nzelu CG, et al. Mitochondria directly interact with the nuclear pore complex. Nature. 18 juin 2026;654(8119):786-97. doi:10.1038/s41586-026-10588-3

47. Sriram K, Qi Z, Yuan D, Malhi NK, Liu X, Calandrelli R, et al. Regulation of nuclear transcription by mitochondrial RNA in endothelial cells. eLife. 22 janv 2024;13:e86204. doi:10.7554/eLife.86204

48. Merlie JP, Sanes JR. Concentration of acetylcholine receptor mRNA in synaptic regions of adult muscle fibres. Nature. sept 1985;317(6032):66–8. doi:10.1038/317066a0

49. Misgeld T, Schwarz TL. Mitostasis in Neurons: Maintaining Mitochondria in an Extended Cellular Architecture. Neuron. nov 2017;96(3):651–66. doi:10.1016/j.neuron.2017.09.055

50. Britto FA, Dumas K, Giorgetti-Peraldi S, Ollendorff V, Favier FB. Is REDD1 a metabolic double agent? Lessons from physiology and pathology. Am J Physiol-Cell Physiol. 1 nov 2020;319(5):C807–24. doi:10.1152/ajpcell.00340.2020

51. Lin K, Wei L, Wang R, Li L, Song S, Wang F, et al. Disrupted methionine cycle triggers muscle atrophy in cancer cachexia through epigenetic regulation of REDD1. Cell Metab. févr 2025;37(2):460–476.e8. doi:10.1016/j.cmet.2024.10.017

52. Pastor F, Dumas K, Barthélémy MA, Regazzetti C, Druelle N, Peraldi P, et al. Implication of REDD1 in the activation of inflammatory pathways. Sci Rep. 1 août 2017;7(1):7023. doi:10.1038/s41598-017-07182-z

53. Williamson DL, Li Z, Tuder RM, Feinstein E, Kimball SR, Dungan CM. Altered nutrient response of mTORC1 as a result of changes in REDD1 expression: effect of obesity vs. REDD1 deficiency. J Appl Physiol. 1 août 2014;117(3):246–56. doi:10.1152/japplphysiol.01350.2013

54. Jimenez-Gallegos D, Corrales WA, Peñaloza-Otarola A, Norambuena-Soto I, Urquiza-Zurich S, Marchese D, et al. Transcriptomic analysis reveals heart failure-related lncRNA LINC00670 as a regulator of colon cancer mesenchymal genes. Biochim Biophys Acta BBA - Mol Basis Dis. nov 2026;1872(8):168389. doi:10.1016/j.bbadis.2026.168389

55. Hain BA, Xu H, VanCleave AM, Gordon BS, Kimball SR, Waning DL. REDD1 deletion attenuates cancer cachexia in mice. J Appl Physiol. 1 déc 2021;131(6):1718–30. doi:10.1152/japplphysiol.00536.2021

