## Supplemental Data for "REDD1 Regulates MERCS, Protein Synthesis and NMJ Stability in Fast Myofibers During Dexamethasone-Induced Muscle Wasting"

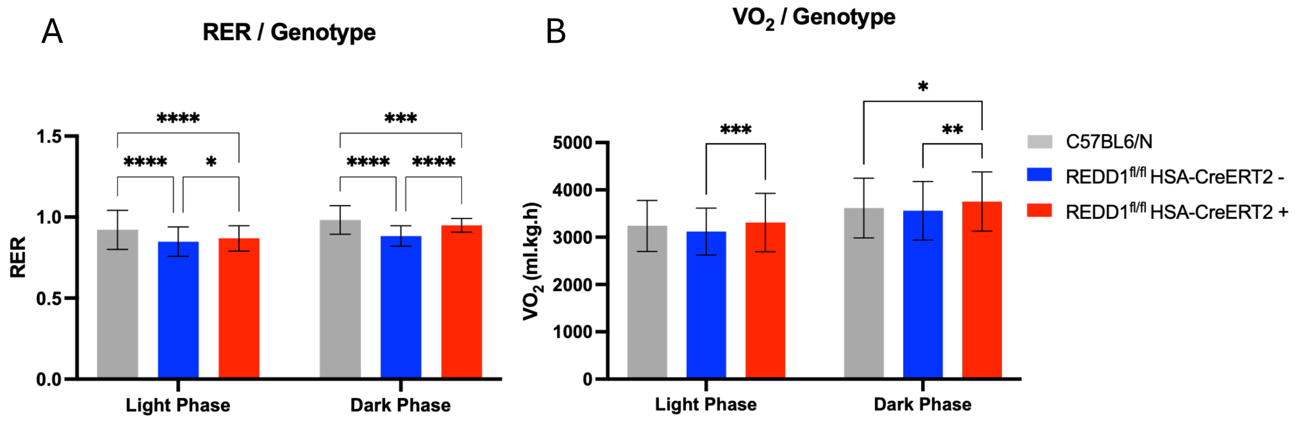
A
RER / Genotype
RER
1.5
1.0
0.5
0.0
****
****
*
***
****
****
Light Phase
Dark Phase
B
VO2 / Genotype
VO2 (ml.kg.h)
5000
4000
3000
2000
1000
0
***
*
**
Light Phase
Dark Phase
C57BL6/N
REDD1fl/fl HSA-CreERT2 -
REDD1fl/fl HSA-CreERT2 +

Figure S1. Whole-body energy metabolism in REDD1 knockout mice

**A.** Respiratory exchange ratio (RER) measured by indirect calorimetry during the light and dark phases in C57BL/6N, REDD1$^{fl/fl}$HSA-CreERT2−(control) and REDD1$^{fl/fl}$HSA-CreERT2+ (KO) mice. RER was calculated as $VCO_2/VO_2$. n=3; Two-way ANOVA, Tukey's post-hoc. **B.** Oxygen consumption ($VO_2$, ml·kg$^{-1}$·h$^{-1}$) during the light and dark phases in the same groups. Two-way ANOVA, Tukey's post-hoc.

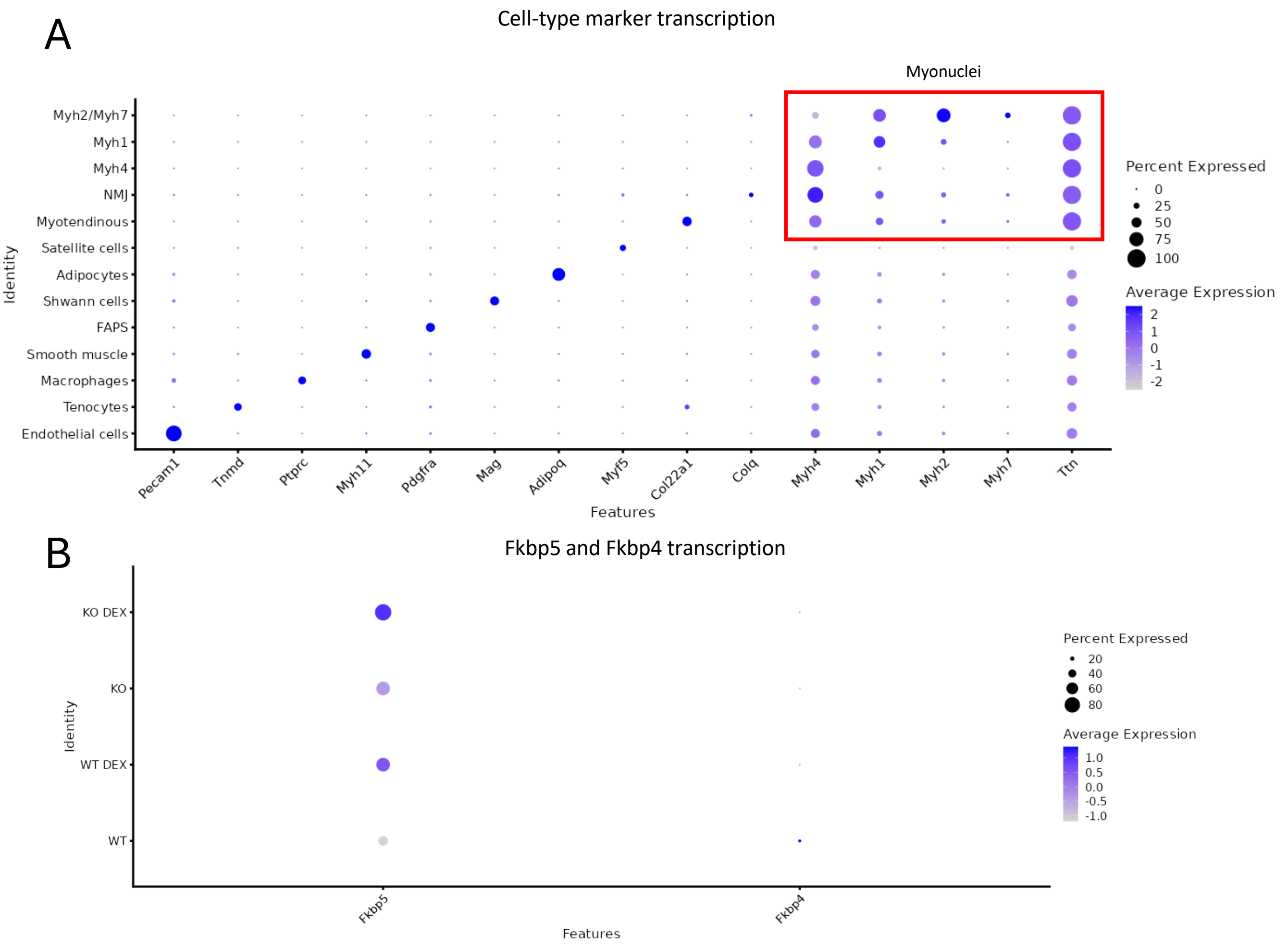
A
Cell-type marker transcription
Myonuclei
Myh2/Myh7
Myh1
Myh4
NMJ
Myotendinous
Satellite cells
Adipocytes
Shwann cells
FAPS
Smooth muscle
Macrophages
Tenocytes
Endothelial cells
Identity
Pecam1
Tnmd
Ptprc
Myh11
Pdgfra
Mag
Adipoq
Myf5
Col22a1
Colq
Myh4
Myh1
Myh2
Myh7
Ttn
Features
Percent Expressed
0
25
50
75
100
Average Expression
2
1
0
-1
-2
B
Fkbp5 and Fkbp4 transcription
KO DEX
KO
WT DEX
WT
Identity
Fkbp5
Fkbp4
Features
Percent Expressed
20
40
60
80
Average Expression
1.0
0.5
0.0
-0.5
-1.0

Figure S2. Cluster annotation and treatment-responsive transcription in myonuclei (snRNA-seq)

**A.** Dot plot of cell-type marker genes used to annotate all nuclei clusters (*Pecam1*, *Tnmd*, *Ptprc*, *Myh11*, *Pdgfra*, *Mag*, *Adipoq*, *Myh5*, *Col22a1*, *Colq*, *Myh4*, *Myh1*, *My h2*, *Myh7*, *Ttn*). Dot size indicates the proportion of expressing nuclei and color intensity indicates normalized expression. **B.** Dot plot of the glucocorticoid-responsive genes *Fkbp5* and *Fkbp4* across WT, WT DEX, KO and KO DEX.

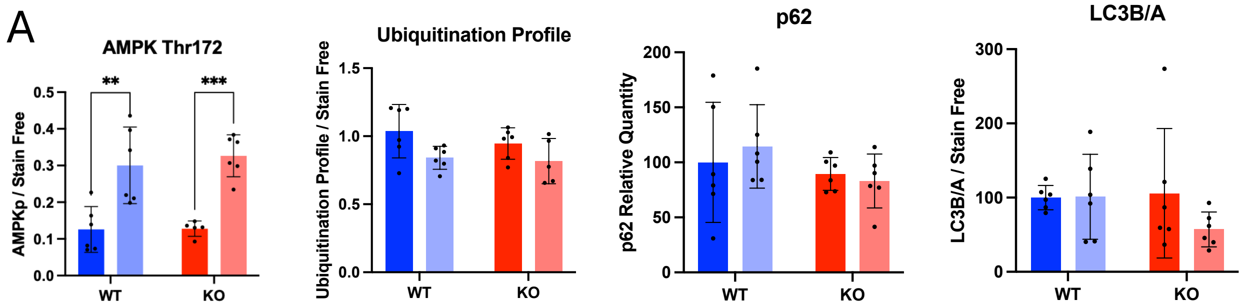
A
AMPK Thr172
AMPKp / Stain Free
0.0
0.1
0.2
0.3
0.4
0.5
**
***
WT
KO
Ubiquitination Profile
Ubiquitination Profile / Stain Free
0.0
0.5
1.0
1.5
WT
KO
p62
p62 Relative Quantity
0
50
100
150
200
WT
KO
LC3B/A
LC3B/A / Stain Free
0
100
200
300
WT
KO

Figure S3. Measurement of protein degradation pathways

**A.** Representative western blots and quantification of phosphorylated AMPK (Thr172), Ubiquitination profile, p62 and LC3B relative to LC3A in quadriceps of WT and KO mice under CTL and DEX. Phosphorylated signal was normalized to Stain-Free. n = 6 per group; two-way ANOVA, genotype × treatment, Tukey's post-hoc.

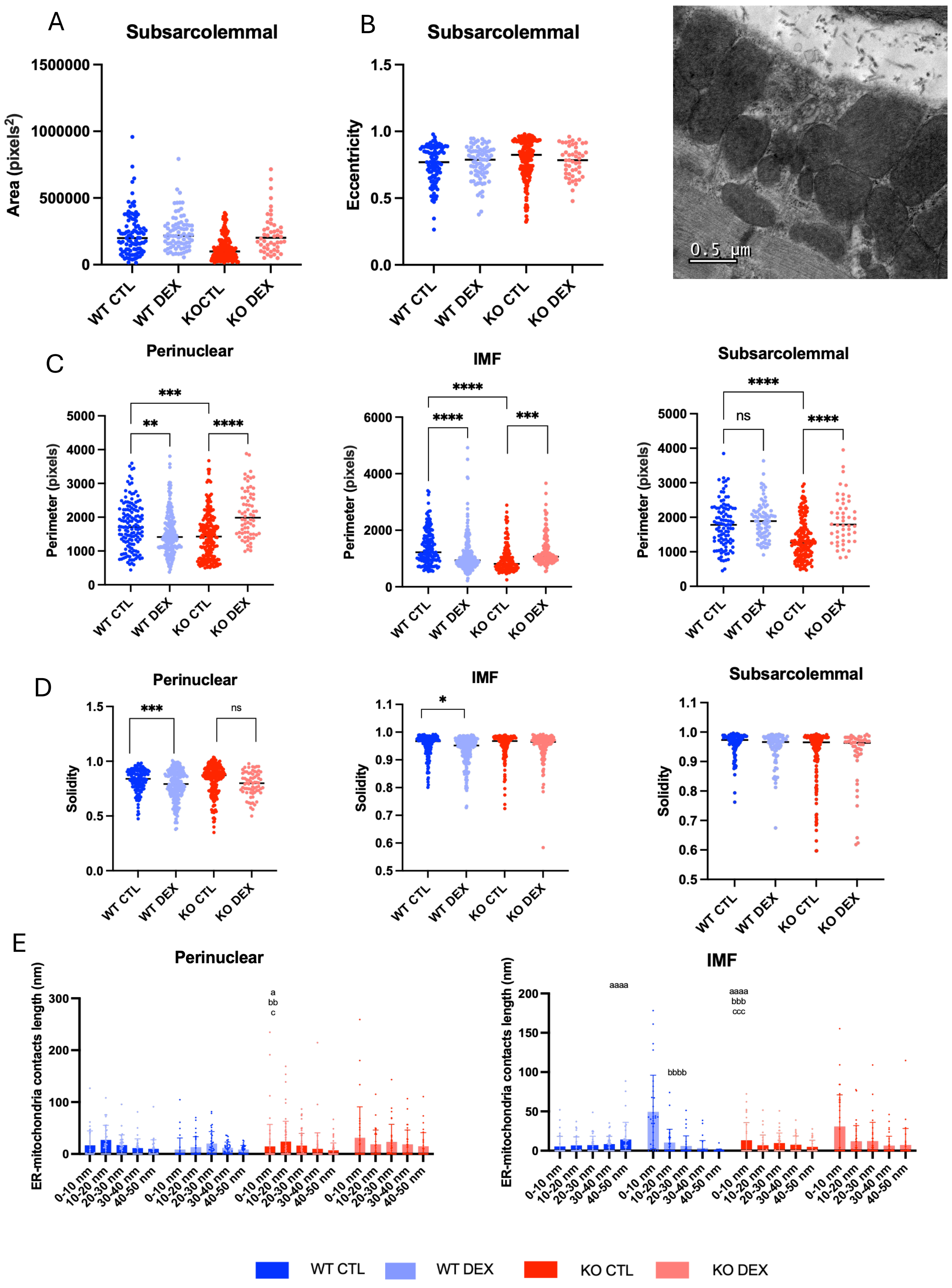
A
Subsarcolemmal
Area (pixels²)
WT CTL
WT DEX
KOCTL
KO DEX
B
Subsarcolemmal
Eccentricity
WT CTL
WT DEX
KO CTL
KO DEX
0.5 μm
C
Perinuclear
Perimeter (pixels)
***
**
****
IMF
****
****
***
Subsarcolemmal
****
ns
****
D
Perinuclear
Solidity
***
ns
IMF
*
Subsarcolemmal
E
Perinuclear
ER-mitochondria contacts length (nm)
a
bb
c
0-10 nm
10-20 nm
20-30 nm
30-40 nm
40-50 nm
IMF
aaaa
bbbb
aaaa
bbb
ccc
WT CTL
WT DEX
KO CTL
KO DEX

Figure S4. Subsarcolemmal mitochondrial morphometry and compartment-wide shape and MERCS metrics

**A.** Subsarcolemmal mitochondrial area (pixels$^2$) across the four groups. **B.** Subsarcolemmal mitochondrial eccentricity across the four groups, with a representative transmission electron micrograph of the subsarcolemmal compartment. **C.** Mitochondrial perimeter (pixels) in the perinuclear, IMF and subsarcolemmal compartments across the four groups. Two-way ANOVA, Tukey's post-hoc. The three compartments were analyzed separately. **D.** Mitochondrial solidity in the same three compartments across the four groups. Two-way ANOVA, Tukey's post-hoc. **E.** Distribution of ER–mitochondria contact length (nm) binned by minimum distance in the perinuclear and IMF compartments across the four groups.

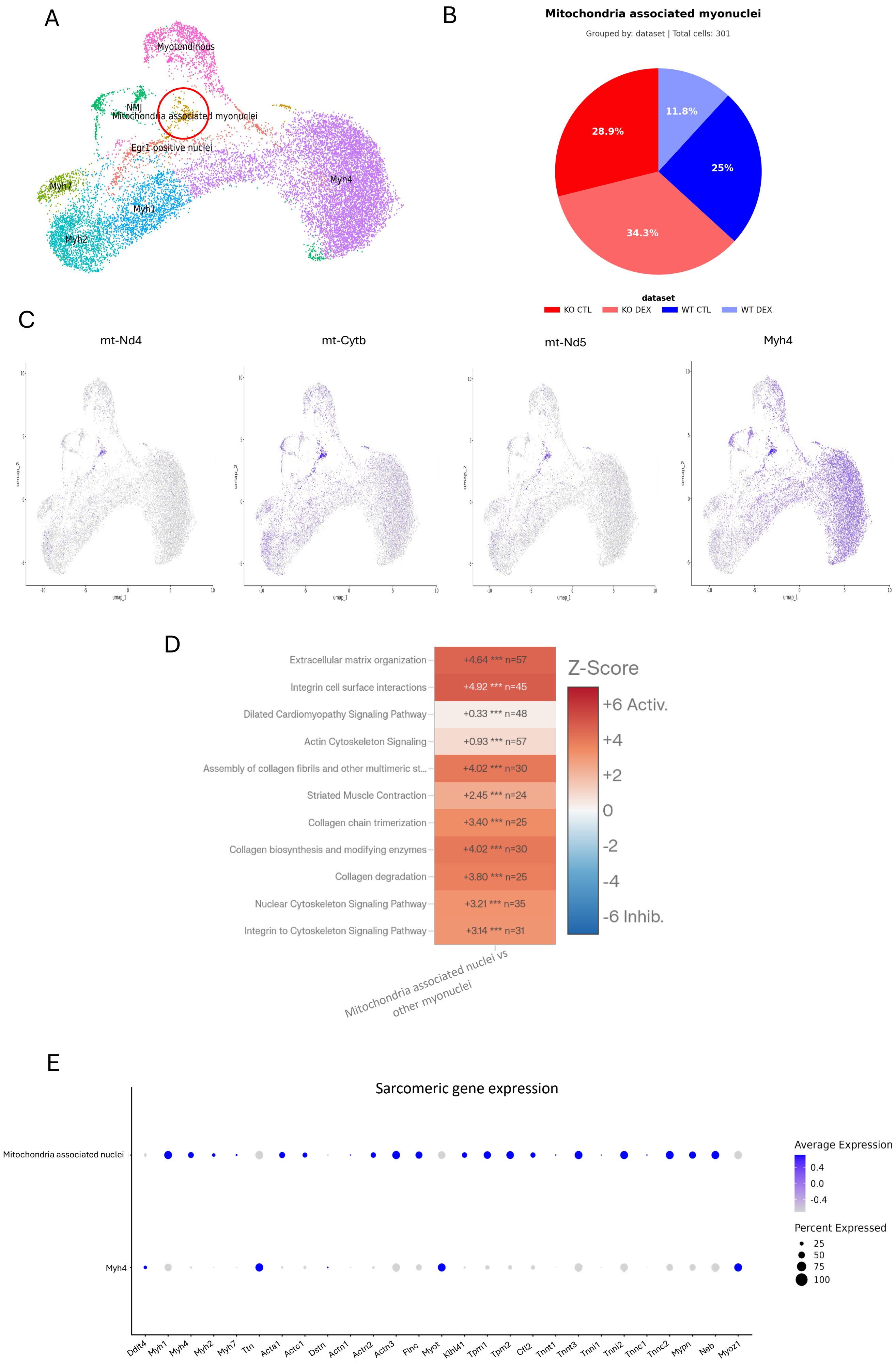
A
Myotendinous
NMJ
Mitochondria associated myonuclei
Egr1 positive nuclei
Myh7
Myh1
Myh2
Myh4
B
Mitochondria associated myonuclei
Grouped by: dataset | Total cells: 301
11.8%
28.9%
25%
34.3%
dataset
KO CTL
KO DEX
WT CTL
WT DEX
C
mt-Nd4
mt-Cytb
mt-Nd5
Myh4
D
Extracellular matrix organization
+4.64 *** n=57
Integrin cell surface interactions
+4.92 *** n=45
Dilated Cardiomyopathy Signaling Pathway
+0.33 *** n=48
Actin Cytoskeleton Signaling
+0.93 *** n=57
Assembly of collagen fibrils and other multimeric st...
+4.02 *** n=30
Striated Muscle Contraction
+2.45 *** n=24
Collagen chain trimerization
+3.40 *** n=25
Collagen biosynthesis and modifying enzymes
+4.02 *** n=30
Collagen degradation
+3.80 *** n=25
Nuclear Cytoskeleton Signaling Pathway
+3.21 *** n=35
Integrin to Cytoskeleton Signaling Pathway
+3.14 *** n=31
Mitochondria associated nuclei vs other myonuclei
Z-Score
+6 Activ.
+4
+2
0
-2
-4
-6 Inhib.
E
Sarcomeric gene expression
Mitochondria associated nuclei
Myh4
Ddit4
Myh1
Myh4
Myh2
Myh7
Ttn
Acta1
Actc1
Dstn
Actn1
Actn2
Actn3
Flnc
Myot
Klhl41
Tpm1
Tpm2
Cfl2
Tnnt1
Tnnt3
Tnni1
Tnni2
Tnnc1
Tnnc2
Mypn
Neb
Myoz1
Average Expression
0.4
0.0
-0.4
Percent Expressed
25
50
75
100

Figure S5. A distinct myonuclear population enriched in mitochondrial transcripts and sarcomeric genes is present across genotypes and glucocorticoid treatment

**A.** UMAP projection of myonuclei (snRNA-seq), with the mitochondria-associated myonuclear cluster circled. Surrounding clusters (Myh4, Myh1, Myh2, Myh7, myotendinous, NMJ, Egr1-positive nuclei) are labelled. **B.** Composition of the mitochondria-associated myonuclear cluster (301 nuclei total) by dataset: WT CTL, WT DEX, KO CTL, KO DEX. **C.** Feature plots of mitochondrially encoded transcripts (*mt-Nd4*, *mt-Cytb*, *mt-Nd5*) and *Myh4* on the myonuclei UMAP. Color intensity indicates normalized transcript expression. **C.** Ingenuity Pathway Analysis (IPA) of genes differentially expressed in mitochondria-associated nuclei versus other myonuclei. Pathways are colored by activation Z-score **E.** Dot plot of sarcomeric gene expression (*Ddit4, Myh4/1/2/7, Ttn, Acta1, Actc1, Dstn, Actn1/2/3, Flnc, Myot, Klhl41, Tpm1/2, Cfl2, Tnnt1/3, Tnni1/2, Tnnc1/2, Mypn, Neb, Myoz1*) in mitochondria-associated nuclei versus Myh4 myonuclei. Dot size indicates the proportion of expressing nuclei and color intensity indicates normalized transcript expression.

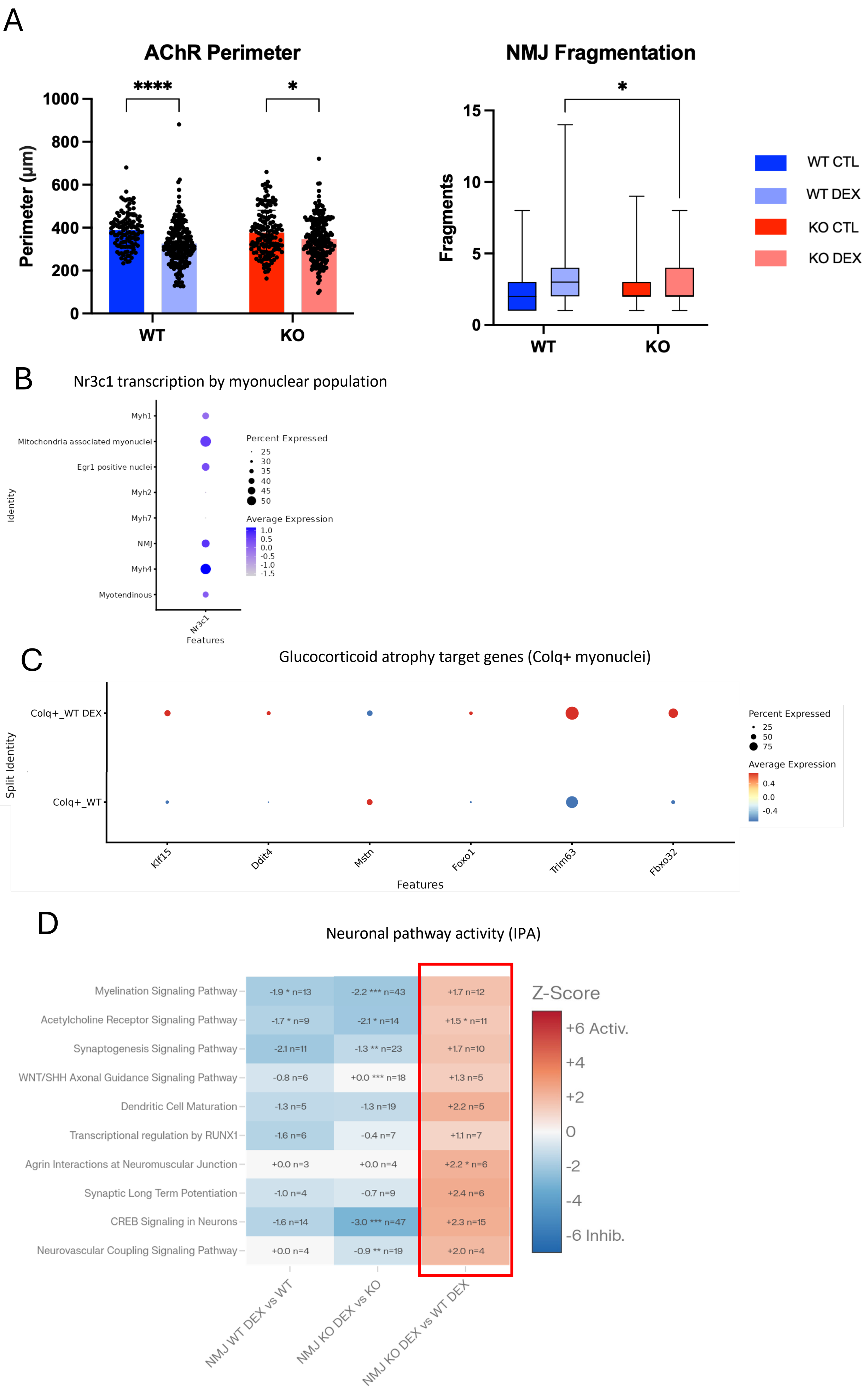
A
AChR Perimeter
Perimeter (µm)
0
200
400
600
800
1000
****
*
WT
KO
NMJ Fragmentation
Fragments
0
5
10
15
*
WT
KO
WT CTL
WT DEX
KO CTL
KO DEX
B
Nr3c1 transcription by myonuclear population
Identity
Myh1
Mitochondria associated myonuclei
Egr1 positive nuclei
Myh2
Myh7
NMJ
Myh4
Myotendinous
Nr3c1
Features
Percent Expressed
25
30
35
40
45
50
Average Expression
1.0
0.5
0.0
-0.5
-1.0
-1.5
C
Glucocorticoid atrophy target genes (Colq+ myonuclei)
Split Identity
Colq+_WT DEX
Colq+_WT
Klf15
Ddit4
Mstn
Foxo1
Trim63
Fbxo32
Features
Percent Expressed
25
50
75
Average Expression
0.4
0.0
-0.4
D
Neuronal pathway activity (IPA)
Myelination Signaling Pathway: -1.9 * n=13; -2.2 *** n=43; +1.7 n=12
Acetylcholine Receptor Signaling Pathway: -1.7 * n=9; -2.1 * n=14; +1.5 * n=11
Synaptogenesis Signaling Pathway: -2.1 n=11; -1.3 ** n=23; +1.7 n=10
WNT/SHH Axonal Guidance Signaling Pathway: -0.8 n=6; +0.0 *** n=18; +1.3 n=5
Dendritic Cell Maturation: -1.3 n=5; -1.3 n=19; +2.2 n=5
Transcriptional regulation by RUNX1: -1.6 n=6; -0.4 n=7; +1.1 n=7
Agrin Interactions at Neuromuscular Junction: +0.0 n=3; +0.0 n=4; +2.2 * n=6
Synaptic Long Term Potentiation: -1.0 n=4; -0.7 n=9; +2.4 n=6
CREB Signaling in Neurons: -1.6 n=14; -3.0 *** n=47; +2.3 n=15
Neurovascular Coupling Signaling Pathway: +0.0 n=4; -0.9 ** n=19; +2.0 n=4
NMJ WT DEX vs WT
NMJ KO DEX vs KO
NMJ KO DEX vs WT DEX
Z-Score
+6 Activ.
+4
+2
0
-2
-4
-6 Inhib.

Figure S6. Glucocorticoid receptor expression, atrophy-gene induction and neuronal pathway activity in synaptic (Colq$^{+}$/NMJ) myonuclei

**A.** Left: acetylcholine receptor (AChR) endplate perimeter (µm) quantified from α-bungarotoxin–stained NMJs in WT and KO muscle under CTL and DEX. Right: NMJ fragmentation, expressed as the number of AChR fragments per endplate. Two-way ANOVA, Tukey's post hoc. **B.** Dot plot of *Nr3c1* (glucocorticoid receptor) expression across myonuclear populations. Dot size indicates the proportion of expressing nuclei and color intensity indicates normalized expression. **C.** Dot plot of glucocorticoid atrophy target genes (*Klf15*, *Ddit4*, *Mstn*, *Foxo1*, *Trim63*, *Fbxo32*) in Colq$^{+}$ myonuclei, WT versus WT DEX. Dot size indicates the proportion of expressing nuclei and color intensity indicates normalized expression. **D.** IPA of neuronal signaling pathways in NMJ myonuclei across three contrasts (NMJ WT DEX vs WT; NMJ KO DEX vs KO; NMJ KO DEX vs WT DEX). Cells report activation Z-score, enrichment significance and overlapping molecule number.

Table 1. List of antibodies used for Western-Blotting

| | | | |
|---|---|---|---|
| REDD1 | Protein Tech #10-638 | 1:500 | WB |
| Puromycin | Millipore #MABE343 | 1:500 | WB |
| Akt Thr308 | Cell Signalling #13038 | 1:1000 | WB |
| Total Akt | Cell Signalling #9272 | 1:1000 | WB |
| S6 Ser 235 | Cell Signalling #4856 | 1:1000 | WB |
| Total S6 | Cell Signalling #2217 | 1:1000 | WB |
| 4EBP1 Thr37/36 | Cell Signalling #2855 | 1:1000 | WB |
| Total 4EBP1 | Cell Signalling #9452 | 1:1000 | WB |
| ULK1 Ser757 | Cell Signalling #6888 | 1:1000 | WB |
| AMPK Thr172 | Cell Signalling #2535 | 1:500 | WB |
| Total AMPK | Cell Signalling #2603 | 1:1000 | WB |
| LC3 | Cell Signalling #3968 | 1:1000 | WB |
| p62 | Cell Signalling #5114 | 1:1000 | WB |
| Ubiquitination profile | Santa Cruz #SC8017 | 1:1000 | WB |
| MYH7 | DSHB BA-F8 | 1:10 | IF |
| MYH2 | DSHB SC-71 | 1:200 | IF |
| MYH4 | DSHB BF-F3 | 1:100 | IF |
| Laminin | Thermo Fisher #L9393 | 1:750 | IF |
| VDAC | Abcam #AB14734 | 1:200 | PLA |
| IP3R | Abcam #AB5804 | 1:200 | PLA |

Table 2. Primers used for RT-qPCR

| | | |
|---|---|---|
| REDD1 | CAGAGAAGAGGGCCTTGA | CCATCCAGGTATGAGGAGTCTT |
| FOXO1 | GCTGGGTGTCAGGCTAAGAG | AGGGCATCTTTGGACTGCT |
| FOXO3 | GGAAATGGGCAAAGCAGA | AAACGGATCACTGTCCACTTG |
| MAFbx | AGTGAGGACCGGCTACTGTG | GATCAAACGCTTGCGAATC |
| MuRF1 | TCCTGCAGAGTGACCAAGG | GGCGTAGAGGGTGTCAAAC |
